# Long-sperm precedence and other cryptic female choices in *Drosophila melanogaster*

**DOI:** 10.1101/2024.04.25.591180

**Authors:** Brooke Peckenpaugh, Joanne Y. Yew, Leonie C. Moyle

## Abstract

Females that mate multiply make postmating choices about which sperm fertilize their eggs (cryptic female choice); however, the male characteristics they use to make such choices remain unclear. In this study, we sought to understand female sperm use patterns by evaluating whether *Drosophila melanogaster* females adjust sperm use (second male paternity) in response to four main factors: male genotype, male courtship effort, male pheromone alteration, and male postmating reproductive morphology. Our experiment was replicated across four different *D. melanogaster* lines, in a full factorial design, including a pheromone manipulation in which second males were perfumed to resemble heterospecific (*D. yakuba*) males. We found that females prefer longer sperm—regardless of mating order—in almost all contexts; this observed pattern of ‘long-sperm precedence’ is consistent with female postmating choice of high-fitness male traits. Nonetheless, we also found that this general preference can be plastically altered by females in response to effects including perfuming treatment; this differential female sperm use is between otherwise identical males, and therefore solely female-mediated. Furthermore, our finding that females exercise choice using diverse criteria suggests a possible mechanism for the maintenance of variation in sexually selected male traits.

**Teaser text:** How do females make choices about which sperm will fertilize their eggs? In this study, we assess four geographically diverse *Drosophila melanogaster* lines to show that females strongly prefer long sperm—a pattern consistent with female preference for high-fitness males. In addition, we perfumed males to show that females can adjust this general preference based on other factors, including pheromones. Because this adjustment occurs between genotypically identical males, it is entirely female-mediated.

## 1 Introduction

Reproduction in nature is costly; organisms are therefore expected to use their reproductive resources judiciously and to adjust them according to reproductive context. Examples of such behavioral reproductive plasticity span diverse traits and taxa, from palm trees (e.g., sex allocation in *Attalea speciosa*, Barot et al. 2005) to guppies (e.g., sperm use in *Poecilia reticulata*, Gasparini & Evans 2018) to flies (e.g., aggression in *Drosophila melanogaster*, Nandy et al. 2016). One specific strategy for plastically allocating reproductive resources is to adjust choices about reproductive partners. Females, in particular, are known to do this in numerous ways prior to mating. For instance, in red jungle fowl, females choose to mate with males with brighter feathers and longer tails; these traits are more prominent in unparasitized males and are therefore honest indicators of male health (Zuk et al. 1990). While many such cases involve choices exercised before mating, some organisms also have the ability to selectively allocate postmating reproductive resources; for instance, female guppies fertilize more eggs when mating with more colorful focal males in comparison to stimulus males (Pilastro et al. 2004). However, because most of these choices are made during and after mating events, it remains hard to differentiate the relative contributions of each mating partner, the criteria on which their choices are based, and therefore the degree to which these choices could be adaptive.

Interestingly, in *Drosophila*, postmating reproductive plasticity has primarily been studied in males. Even under optimal conditions, *D. melanogaster* males are known to be ejaculate-limited, with their progeny output declining after two to five matings in a day (Douglas et al. 2020, Hopkins et al. 2019, Linklater et al. 2007). Plastic allocation of reproductive resources could therefore allow them to prioritize sperm and seminal fluid for preferred mates. Consistent with this, *D. melanogaster* males use premating information, such as female condition and the relative abundance of potential sexual competitors, to modulate ejaculate composition (Garbaczewska et al. 2013, Lüpold et al. 2011, Sirot et al. 2011, Sirot 2019, Wigby et al. 2009). *D. melanogaster* males also allocate more resources to matings with large females—an investment that reduces their success at defensive sperm competition in subsequent matings (Anastasio et al. 2023). Together, these studies broadly indicate that costs to *D. melanogaster* males drive plasticity in their reproductive allocation during mating, as well as identify specific criteria used in making these behavioral decisions.

Like males, females are also expected to selectively allocate reproductive resources (i.e., eggs) at postmating stages, through the action of “cryptic female choice”—the female-mediated physical and chemical factors that determine which sperm are used to fertilize eggs after mating (Eberhard 1996, Firman et al. 2017, Thornhill 1983). In *Drosophila,* several proximate mechanisms of cryptic female choice involve modulating the storage of sperm prior to fertilization. *Drosophila* females have two types of sperm storage organs and can bias sperm use through differential sperm storage and/or by preferentially drawing sperm from these alternative organs (Manier et al. 2013b). Females can also manipulate the identity of sperm available in storage by adjusting the time until sperm ejection after specific matings, which terminates the displacement of resident sperm by incoming sperm (Laturney et al. 2018, Lüpold et al. 2013, Manier et al. 2010, Manier et al. 2013a, Snook & Hosken 2004). Finally, it has been proposed that the size of sperm storage organs themselves might provide a mechanism by which females make these choices. For example, females experimentally evolved to have longer seminal receptacles have been shown to select for males with longer sperm (Miller & Pitnick 2002). Even when the specific mechanism is unknown, it is evident that *Drosophila* females can change sperm use patterns depending upon context, including according to male genotype identity (Chow et al. 2010). What is comparatively poorly understood, however, is the evolutionary explanation for these apparent female choices, including the criteria used by females to exercise them.

One reason for this is the challenge of differentiating female choice from primarily male-mediated factors that can also influence the outcome of postmating processes (Firman et al. 2017). For instance, *Drosophila simulans* males that are more attractive to females also sire more offspring as second males; however, it is unclear whether this is because females preferentially use sperm from attractive males, or because attractive males are intrinsically better sperm competitors (Hosken et al. 2008). While the former explanation would be consistent with an adaptive function for cryptic female choice, the latter explanation implies passive female responses to the outcome of male-male sexual selection. One strategy for determining the specific contribution of female choices to patterns of biased sperm use is to evaluate how these patterns change when female perception of male identity is manipulated, without influencing male identity itself. Although female assessment of male identity is likely to be complex, in *Drosophila,* female premating preference primarily depends on auditory and chemosensory cues (Greenspan & Ferveur 2000, Yamamoto & Nakano 1999); manipulating these cues could therefore modify this assessment. Contact pheromones are one such cue. In insects broadly, and flies specifically, these pheromones include cuticular hydrocarbons (CHCs)—waxy compounds on the outer surface (cuticle) of many insects that can be used as sexual pheromones in mate choice by males (Billeter et al. 2009) and females (Scott 1994), both within and between species (Blows and Allan 1998). CHC composition can be artificially altered to assess its effect on mate choice by “perfuming” focal males with the CHCs of other male genotypes (Dyer et. al. 2014, Wang et al. 2011). For instance, *D. melanogaster* males have divergent pheromone profiles from the closely related species *Drosophila yakuba,* both in the abundance of CHCs and in the presence or absence of some CHCs (Figure S1, Khallaf et al. 2021). Perfuming *D. melanogaster* males with CHCs from *D. yakuba* males, immediately prior to mating, can therefore manipulate female perception of male identity while keeping male postmating quality the same.

In this study, we sought to understand the nature and relative importance of factors that females use when exercising cryptic female choice. To do so, we evaluated whether *Drosophila melanogaster* females adjust their sperm use patterns in response to four main factors: male genotype, male courtship effort, male pheromone alteration, and male postmating reproductive morphology (the morphological traits relevant to postmating reproductive success). We manipulated male pheromones via “perfuming” with heterospecific (*D. yakuba*) males—thereby changing the perceived, but not actual, identity of these treatment males. We also evaluated postmating reproductive trait variation in males and females (testis length, sperm length, male reproductive tract [MRT] mass, sperm storage organ size, female reproductive tract [FRT] mass) to assess their contribution to sperm preference patterns. We assessed how second-male paternity (proportion of offspring sired) changed according to each factor, by mating males of each type to females that had previously mated with a consistent first-male tester genotype. Our experiment was replicated across four different *D. melanogaster* lines in a full factorial design, to evaluate whether the effects of these factors on sperm use depend upon female genotype, male genotype, or their interaction. We find clear evidence that females adjust their use of sperm from the second male. In particular, females show a general preference for longer sperm, whether that sperm is from the first or the second male—i.e. ‘long-sperm precedence’ rather than ‘last-male sperm precedence’ (Laturney et al. 2018). Nonetheless, we also find that this general preference can be overridden in specific contexts by female-mediated changes in response to male genotype, courtship effort, and perfuming treatment. The presence and direction of these cryptic choices vary in complex ways, including evidence that females of some genotypes adjust sperm use based on their perception of male identity, regardless of actual identity. These data allow us to draw broader inferences about the possible criteria for, as well as the evolutionary causes and consequences of, such choices.

## 2 Methods

### 2.1 Fly stocks

All stocks were reared on standard media prepared by the Bloomington Drosophila Stock Center and kept at room temperature (∼22°C). For this experiment we used four lines from among the 15 *D. melanogaster* founder lines in the Drosophila Synthetic Population Resource (DSPR) (King et al. 2012). These fully-sequenced founder lines represent a diverse collection of genotypes from different locations around the world, and are known to vary in reproductive traits (i.e., male courtship song, see Pischedda et al. 2014a). Our four DSPR founder lines (A5 [Greece], A7 [Taiwan], B4 [California], and B7 [Malaysia]) were used as focal genotypes for females and second males in each manipulation. The GFP-labeled *D. melanogaster* line (stock #32170 from the Bloomington *Drosophila* Stock Center) was used as the first tester male. The *D. yakuba* line yak_S09_L38 was the heterospecific donor male for the heterospecific-perfumed treatment (collected by B. Cooper in Sao Tome in 2009). For each conspecific-perfumed treatment, target males were perfumed (see methods below) with males from their own genotype.

### 2.2 Experimental design

To assess patterns of preferential sperm use, including after remating with conspecific- vs. heterospecific-perfumed males, we mated *D. melanogaster* females sequentially to two males: first, to *D. melanogaster* GFP-labeled tester males, and second, to *D. melanogaster* target males. Using a GFP-labeled first male allowed us to assess paternity share (first [tester] versus second [target] males) of offspring from each female) by examining the phenotypes of offspring generated after the second mating. Any offspring not expressing GFP are sired by the second focal male. For our CHC manipulation, the second, target males were either perfumed by genotypically identical males (conspecific-perfumed control) or perfumed by heterospecific *D. yakuba* males (Figure 1).

**Figure 1.**
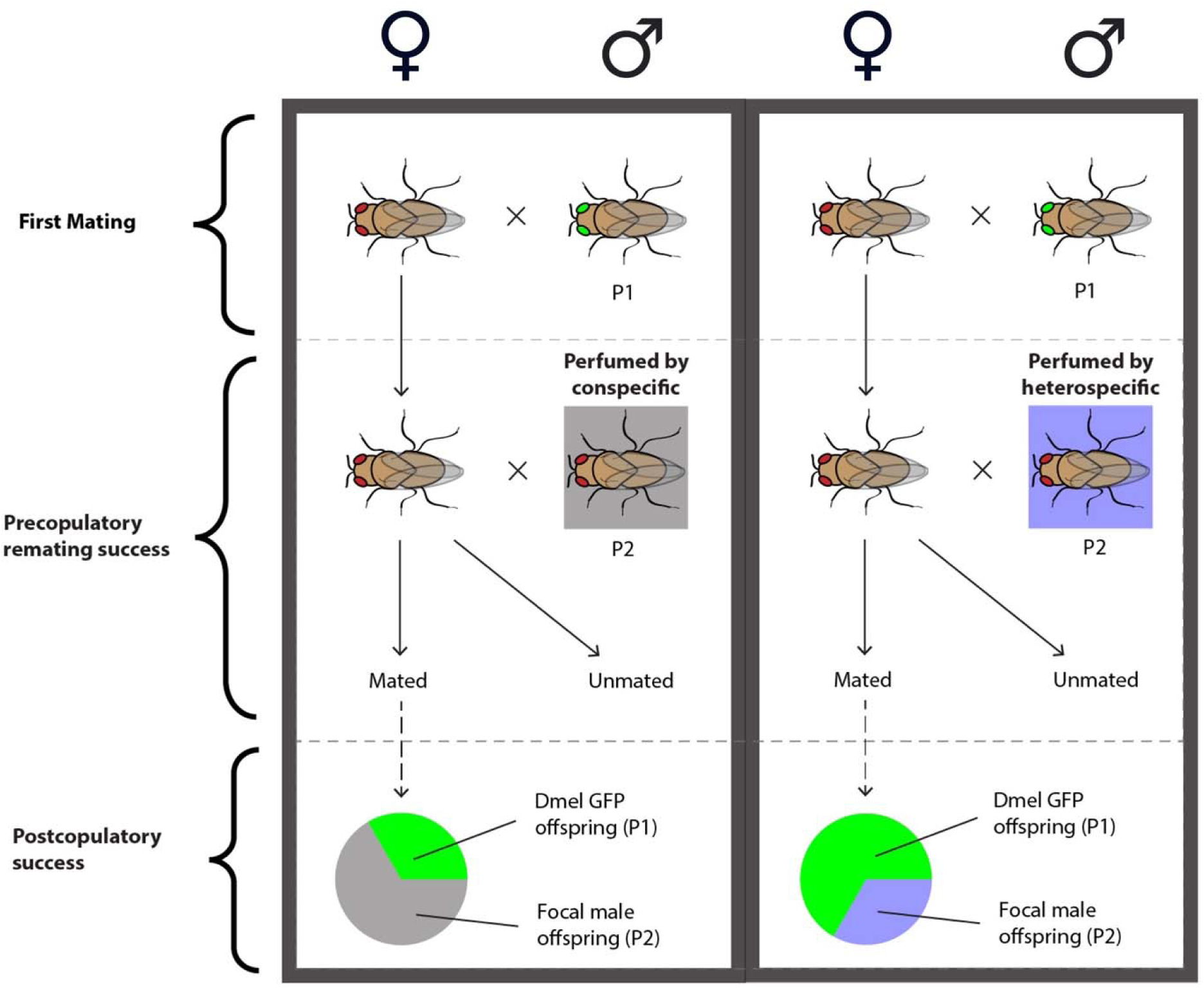
Experimental procedure for assessing paternity share of second (focal) males. Females were first mated with a GFP tester male. After 48 hours, these females were paired for second matings with focal males, which were either perfumed by conspecifics (left, control) or by heterospecifics (right, treatment). Unmated females were removed from the experiment. In remated females, we assessed paternity for first and second males based on the presence/absence of the GFP marker in resulting offspring. **Alt text:** A diagram of our experimental design. Females are mated to two males before their offspring are assessed, such that we can compare patterns of paternity between male genotypes and perfuming treatments.

We replicated this experimental procedure across all four *D. melanogaster* genotypes, in a full factorial design. For each line, we set up matings between females and second males from each of the four DSPR lines, including males from the females’ own genotype. This created 16 cross types in total. For each cross-type, we assessed paternity share from crosses with conspecific-perfumed males and with heterospecific-perfumed males. Each of these cross types was repeated until successfully replicated at least five times, resulting in over 160 total crosses.

### 2.3 Perfuming manipulation

Males used in the second mating were perfumed in the morning before setting up crosses. To do so, we vortexed four target males with 30 donor males, either of the same *D. melanogaster* genotype (for conspecific-perfumed control crosses) or with 30 heterospecific (*D. yakuba*) donor males (for heterospecific-perfumed treatment crosses). For both treatments, we first removed wings from the donor males under CO_2_ anesthesia in order to differentiate them from target males. The flies were vortexed on medium strength for one minute total, in three 20-second increments with 20-second breaks in between (Pischedda et al. 2014b). The target males were separated immediately after vortexing and isolated individually in vials. They were allowed to recover for one hour before setting up crosses.

### 2.4 Paternity assay

*D. melanogaster* males were isolated and aged for seven days, while females were isolated and aged for two days before setting up the first cross with a GFP-labeled tester male. We allowed mating for 24 hours before removing the males. We allowed the isolated female to lay eggs for an additional 24 hours and verified successful matings by observing larval activity. Unmated females (those that failed to produce larvae after the first mating) were removed from the experiment. Mated females were then moved to a new vial to be paired with the second male. To determine whether and the degree to which females exhibit premating discrimination against heterospecific-perfumed males, we observed a subset of crosses for two hours to observe the number of second matings within this period and recorded the time until remating (latency) and copulation duration for each (these data are not presented because only 9 of 162 scored pairings resulted in copulation within two hours). Regardless of whether matings were observed in the first two hours, seven-day-old second males were housed with females for 24 hours. Males were then removed and the females were transferred to a new vial. Females continued to be transferred to new vials every 24 hours for three days. Progeny in these vials were allowed to mature into adulthood, then counted and assessed for GFP to determine paternity (see details below). All cross types were repeated until we had five successful replicates (crosses in which there is evidence for both a first and second mating). Conspecific- and heterospecific-perfumed treatment crosses for each genotype were completed in parallel and in a randomized order, and the researcher counting progeny was blind to which cross type or treatment they were measuring.

Upon eclosion, all progeny from the three post-remating vials were anesthetized with CO_2_ and viewed under a Leica M205FA stereo scope equipped with a UV light for visualizing GFP. The presence or absence of GFP in the ocelli of the eye were used as a marker of paternity; offspring with GFP ocelli were sired by the first male and were thus counted as P1. Progeny with wild-type ocelli were sired by the second male and were counted as P2. If no wild-type progeny were present in the three post-remating vials, we assumed that no remating event occurred, and these were scored as “unremated”.

### 2.5 Cuticular hydrocarbon extraction and quantification

We confirmed the efficacy of perfuming using gas chromatography–mass spectrometry (GC/MS) analysis. We extracted cuticular hydrocarbons from males of all four genotypes, for both conspecific-perfumed and heterospecific-perfumed treatments. Following treatment of seven-day old virgin males, cuticular hydrocarbons were extracted from pooled samples by placing five males in a 1.8 mL glass vial (Wheaton 224740 E-C Clear Glass Sample Vials) with 120 μL of hexane (Sigma Aldrich, St Louis, MO, USA) spiked with 10 μg/mL of hexacosane (Sigma Aldrich). Hexane is a standard non-polar solvent used to extract hydrophobic molecules from insect cuticles. The alkane hexacosane is not endogenously produced by *Drosophila* and serves as an internal standard. After 20 min, 100 μL of the solution was removed to a sterilized 1.8 mL glass vial (Wheaton 224740 E-C Clear Glass Sample Vials) and allowed to evaporate under a fume hood, such that only the extracted cuticular hydrocarbons remained at the bottom of the tube. Extracts were then stored at −20°C until analysis. The samples were reconstituted in 100 μL of hexane immediately before analysis. Five replicate samples, consisting of five flies per sample (25 flies total), were prepared for each sample type.

We performed gas chromatography–mass spectrometry (GC/MS) analysis on a 7820A GC system equipped with a 5975 Mass Selective Detector (Agilent Technologies, Inc., Santa Clara, CA, USA) and a HP-5ms column ((5%-Phenyl)-methylpolysiloxane, 30 m length, 250 μm ID, 0.25 μm film thickness; Agilent Technologies, Inc.). Electron ionization (EI) energy was set at 70 eV. One microliter of the sample was injected in splitless mode and analyzed with helium flow at 1 mL/min. The column was set at 40°C and held for 3 min, increased to 200°C at a rate of 35°C/min, then increased to 280°C at a rate of 3°C/min and held for 15 min. Chromatograms and spectra were analyzed using MSD ChemStation (Agilent Technologies, Inc.). We identified CHCs based on retention time and electron ionization fragmentation pattern. The abundance of each compound was then quantified by normalizing the area under each CHC peak to the area of the hexacosane signal using homebuilt peak selection software (personal correspondence, Dr. Scott Pletcher, Univ. of Michigan).

### 2.6 Postmating reproductive trait morphology

We measured several elements of reproductive trait morphology in each of our lines, and in the tester line, in order to assess their potential influence on patterns of female preferential sperm use.

#### 2.6.1 Thorax length, testis length, SR length, and ST area

We collected both female and male virgins and aged them to reproductive maturity (N=5 replicates per sex per genotype). We measured thorax length for all flies; this is a standard measure of body size in *Drosophila* studies because it strongly correlates with the size of other characters such as wing length (Robertson & Reeve 1952). We then dissected the reproductive tract into Grace’s Insect Medium. For females, we separated the seminal receptacle (SR) and a randomly chosen spermatheca (ST) from the rest of the reproductive tract and transferred them to a slide. For males, a randomly chosen testis was separated from the rest of the reproductive tract and transferred to a slide. In both cases, we then applied a cover slip and took images under an EVOS FL digital inverted microscope. Using ImageJ, we measured the length of the SR, the height and width of the ST, and the length of the testis, from these images. Measurements were replicated across five individuals per genotype.

#### 2.6.2 Sperm length

We dissected sperm out of the seminal vesicles of reproductively mature virgin males into Grace’s insect medium on a slide (N=5 replicates per genotype). Sperm were gently teased apart with a probe when necessary. The slides were stained with SpermBlue (Microptic), and imaged under an EVOS FL digital inverted microscope. In ImageJ, we measure sperm length for 5-10 randomly chosen sperm per individual.

#### 2.6.3 Body mass and reproductive tract mass

We collected both female and male virgins and aged them to reproductive maturity (N=5 replicates per sex, per genotype). We measured body and reproductive tract mass according to Holman et al. (2008). Flies were dissected in distilled water onto a piece of preweighed foil. For females, the reproductive tract (including the bursa, SR, ST, parovaria, and ovaries) was removed and transferred to another piece of pre-weighed foil. For males, the reproductive tract (including testes, seminal vesicles, accessory glands, and the ejaculatory bulb) was removed and transferred to another piece of pre-weighed foil. For both males and females, the foils holding the body and reproductive systems of the flies were then placed in a drying oven at 60°C for 18 ± 1 h and weighed on a Mettler Toledo UMX2 balance to the nearest 0.1 μg.

### 2.7 Second-male courtship behavior assay

To identify variation in male courtship across genotypes, and to determine whether our experimental manipulation affects male courtship behavior, we recorded male courtship for each cross type on video. We recorded videos using a modified FlyPi setup, which combines a Raspberry Pi (Raspberry Pi Foundation, Cambridge, UK), pi camera, and 3D printed parts (Maia Chagas et al. 2017). Crosses were set up in a modified cell culture plate consisting of six culture wells, each with a 3 cm diameter and a small amount of cornmeal media in the bottom, allowing us to record six crosses simultaneously. Male and female flies were aspirated without anesthetization into each cell culture well, and the plate was videotaped for two hours. The full videos were scored for successful copulations, latency to copulation, and copulation duration. The first 30 minutes of each video was also scored in more detail for the frequency of courtship song, male tapping, number of male licking attempts, and copulation attempts. Recorded matings were set up until we had five replicate recordings per cross type.

### 2.8 Statistical analyses

All statistical analysis and figure construction in this study was performed with R version 4.3.1. All linear models were run using deviation-coded contrasts. This does not change the fit of the model, but affects the interpretation of regression coefficients such that the intercept corresponds to the grand mean and main effect estimates represent deviations from the mean. Model estimates are therefore robust to whichever line is used as the baseline in each comparison, which is preferable in our experiments where none of our genotypes were *a priori* designated as the baseline level.

#### 2.8.1 CHC composition

We assessed variation in CHC composition using a principal components analysis (PCA), which we computed using the prcomp function in R. To test for significant differences among our principal components, we performed a multivariate analysis of variance (MANOVA) using the manova function.

#### 2.8.2 Postmating reproductive trait morphology

We assessed variation in female and male reproductive morphology by fitting linear models using the lm function in R. To test whether male or female genotype predicts the variation we observed in reproductive morphology, we ran linear models for each morphological trait separately as the dependent variable. Male or female genotype was always included as the predictor. The GFP-labelled *D. melanogaster* line was incorporated into each model as a baseline.

#### 2.8.3 Second-male courtship effort, female second-mating frequency, and second-male paternity

We assessed variation in second-male courtship by fitting a linear model using the lm function in R. Second male courtship effort was the dependent variable, with female genotype, male genotype, their interaction, and male perfuming treatment, as predictors.

We assessed variation in female second-mating frequency by fitting a beta-binomial generalized linear model using the betabin function in R. We coded our response as a binomial variable, with one column representing the number of successful second-matings, and the other column representing the number of failed second-matings for each cross type (N=32 total cross types). The model included female genotype, male genotype, and male perfuming treatment as predictors (this dataset did not allow us sufficient power to assess interaction terms).

Variation in second male siring success was also assessed with a beta-binomial generalized linear model. We coded our response as a binomial variable, with one column representing the number of offspring sired by the second male (“successes”), and the other column representing the number of offspring sired by the first male (“failures”) for each cross (N=196 total crosses). Our model included male genotype, female genotype, male perfuming treatment, and all possible interactions (female genotype by male genotype, female genotype by male perfuming treatment, male genotype by male perfuming treatment, and female genotype by male genotype by male perfuming treatment) as predictors. California males, California females, and conspecific-perfumed males were always incorporated into the models as baselines.

#### 2.8.4 Correlations among datasets

Finally, we looked for correlations among our datasets. Across all pairwise genotype combinations we assessed the strength of association between: second-male courtship effort and second-male mating success; second male courtship effort and paternity share (P2); female readiness to remate and second-male paternity share (P2); and, male and female reproductive trait variation and second-male paternity share (P2). To do this, in each case, we computed Spearman’s rank correlation coefficients using the cor.test function in R. Each correlation was assessed separately for experiments involving conspecific-perfumed males and heterospecific-perfumed males.

## 3 Results

### 3.1 Males differ in CHC composition based on genotype and perfuming status

We quantified CHC composition across male genotypes and perfuming treatments to characterize the variation present across lines, as well as to verify that our perfuming treatment was effective. We found that CHC composition varied substantially across male genotypes ((Figure 2a,b, Table S1, Table S2). For each of the first three PCs—which together account for 92.3% of the variation observed—we detected a significant genotype effect (MANOVA *p* < 0.001; Table S1). CHC composition was also shifted by our perfuming treatment, as shown by significant treatment effects for PC3 (*p* = 0.004) and PC4 (*p* < 0.001) (MANOVA; Figure 2b, Table S1). These two PCs account for 8.6% of the variation in CHC composition.

**Figure 2.**
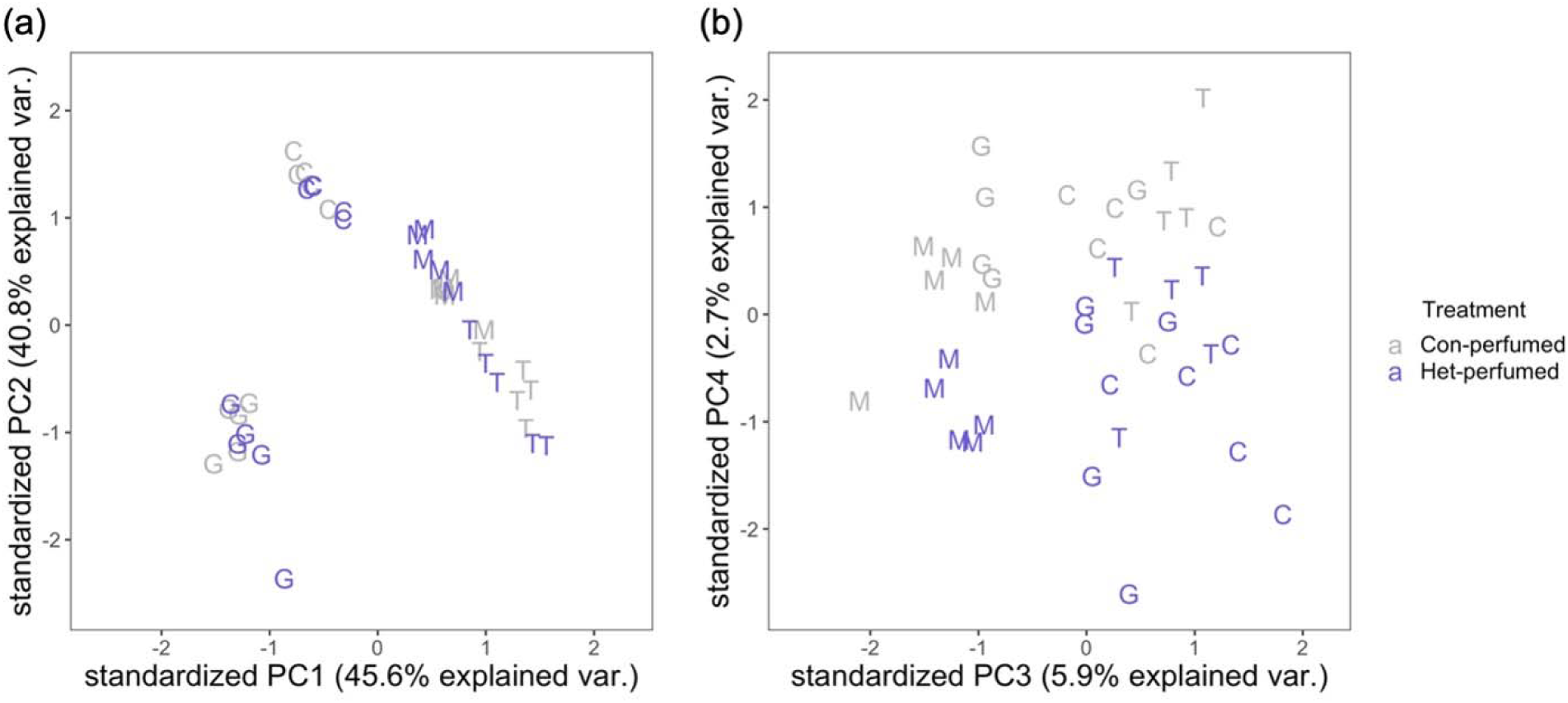
CHC composition varied based on genotype (G, Greece; T, Taiwan; C, California; and M, Malaysia) and perfuming status (conspecific-perfumed males shown in grey, heterospecific-perfumed males shown in purple) (Table S1). Both panels show standardized principal component (PC) values generated from multivariate CHC composition data for replicate males of each genotype and treatment. Left panel: PCs 1 and 2 largely reflect differences between male genotypes. Right panel: PCs 3 and 4 reflect additional shifts in CHC composition associated with heterospecific perfuming treatment. **Alt text:** A principal component analysis of CHC composition. On the left, we see CHC composition clustering according to male genotype. On the right, CHC composition separates out according to perfuming treatment. Points are labeled using the first letter of their genotype.

### 3.2 Postmating reproductive traits differ more among males than among females

We quantified variation in male and female traits, focusing on postmating reproductive traits, to assess whether these might influence patterns of sperm use. We found that all measured male traits differed among genotypes, including two overall size traits and three reproductive size traits (Figure 3, Table S3). Differences among female lines were more modest and only observed for overall size and one reproductive size trait, RT mass (Figure 3, Table S3).

**Figure 3.**
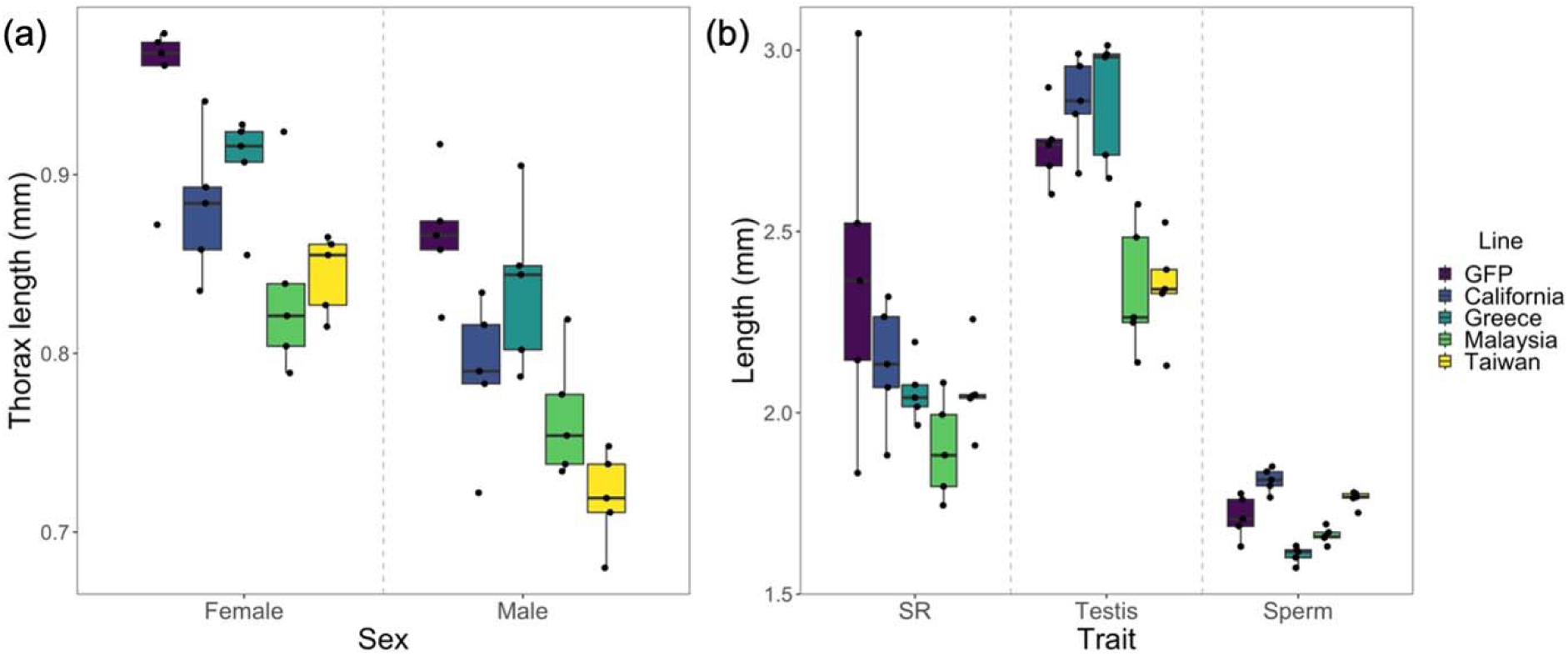
Variation in (a) thorax length in mm and (b) reproductive trait length in mm in females and males across *D. melanogaster* lines. We observed significant variation among lines for thorax length (both males and females), testis length, and sperm length, but SR length did not significantly differ between lines (Table S3). Box plots show the median (central line) and first and third quartiles; whiskers extend to the smallest and largest values (not including outliers >1.5x the inter-quartile range). **Alt text:** A graphical representation of variation in morphology across male and female *D. melanogaster* lines. Data are represented using box plots and individual data points.

In terms of non-reproductive traits, both male and female Taiwan individuals consistently had shorter thoraxes but also the largest dry non-RT body mass. Other differences among lines were sex-specific and often more modest (for example, Greece males had longer thoraxes but reduced dry non-RT body mass, and Malaysia females were smaller than average by both measures) (Table S3).

For reproductive traits, California females exhibited the highest RT dry mass and Taiwan females exhibited the lowest; however, female lines did not differ for either SR length or ST area (Figure S2). In contrast, male lines differed in RT mass (Taiwan males had lower RT mass, while Greece males had higher RT mass, than average), as well as testis length and sperm length (Figure 3, Figure S2). California and Greece males had longer testes than average, while Malaysia and Taiwan males had shorter testes. Interestingly, California and Taiwan males had longer sperm than all other lines (1.814 mm and 1.763 mm on average, respectively; Table S3), and Malaysia and Greece males had significantly shorter sperm (1.662 mm and 1.609 mm on average, respectively; Table S3). GFP sperm length was intermediate (1.713 mm on average; Table S3). This indicates that sperm length and testis length are not closely associated among our *D. melanogaster* lines: Taiwan males had short testes but also had among the longest sperm, while the inverse pattern was observed for Greece males.

### 3.3 Courtship effort varies consistently among male genotypes, but not among female genotypes being courted or with perfuming

We evaluated courtship across each male-female genotype combination to better understand if and/or how premating processes influence cryptic female choices, including via our perfuming treatments. We measured courtship effort as the number of courtship behaviors exhibited per minute until copulation occurred or until our observation period ended (at two hours). Note that only 9 of 162 scored pairings resulted in copulation within two hours. Courtship intensity varied significantly among male genotypes, with Greece and Taiwan males courting significantly less than the others (Figure 4, Table S4). Male courtship intensity was also marginally reduced in heterospecific-perfumed males, but this effect was not statistically significant (Figure 4, Table S4). In addition, although female line identity had only a marginal influence on courtship, we did find that Taiwan females were consistently courted less than average across male genotypes (Figure 4, Table S4), possibly as a result of their smaller size (Turiegano et al. 2012, Table S3). We did not detect any significant female-by-male genotype interactions.

**Figure 4.**
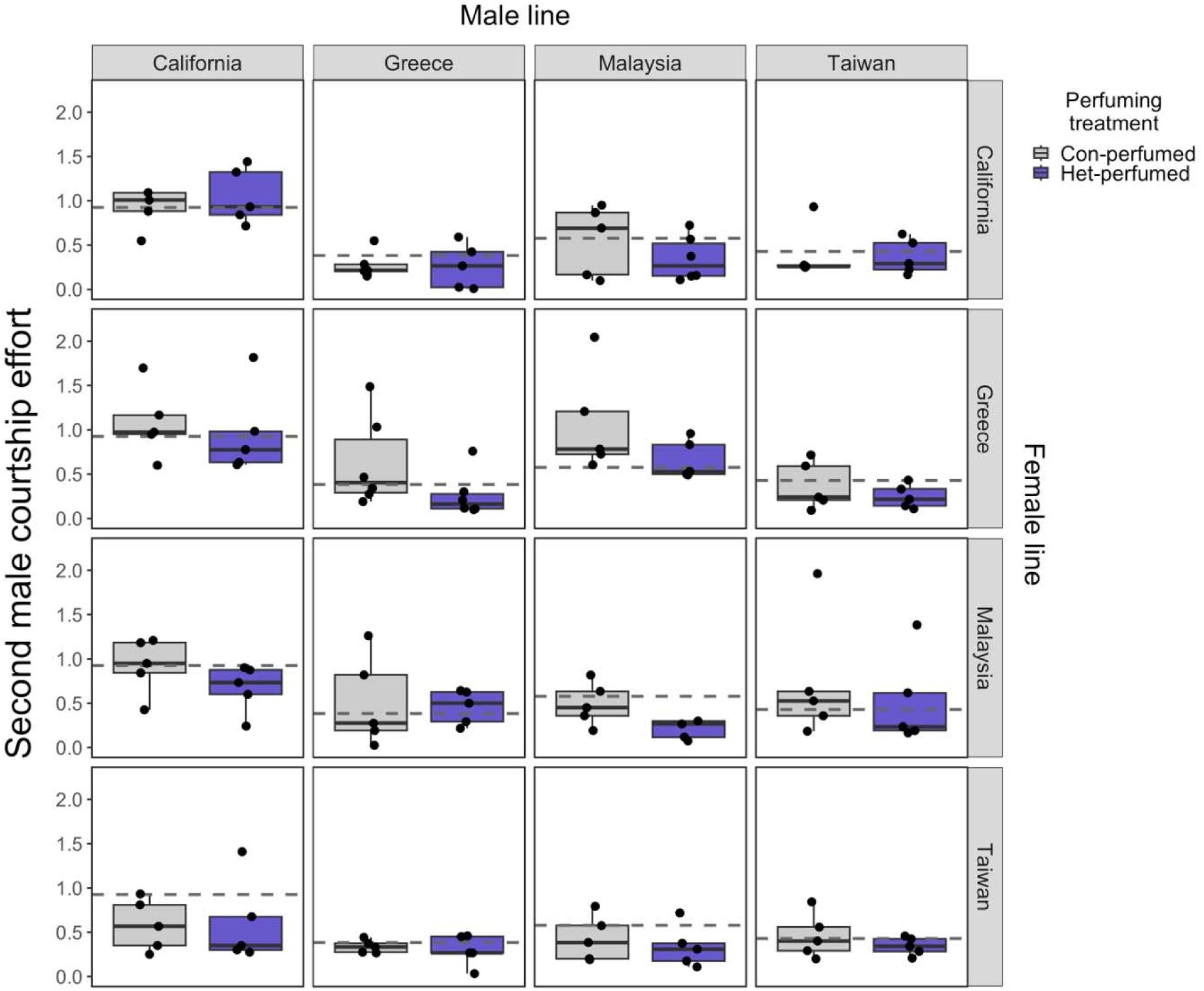
Courtship effort (courtship behaviors per minute) consistently differed between second male genotypes but did not consistently vary by the female genotype being courted or by male perfuming status (Table S4). Conspecific-perfumed males are shown in grey, and heterospecific-perfumed males are shown in purple. The dashed line indicates the average courtship effort for each male line. Box plots show the median (central line) and first and third quartiles; whiskers extend to the smallest and largest values (not including outliers >1.5x the inter-quartile range). The y-axis has been truncated to make variation in the data easier to visualize; the non-truncated data (as used in statistical analyses) is shown in Figure S3. **Alt text:** A matrix of graphs representing second male courtship effort across all possible male by female combinations, divided according to conspecific- or heterospecific-perfumed treatments. Data are represented using box plots and individual data points.

### 3.4 Readiness to remate differs among female genotypes and depends on male genotype, but not on male perfuming status

For each cross type, we assessed whether females chose to mate with second males versus refused the second-male mating. Remating was evidenced by whether a female produced offspring from the second male. We found that female genotypes varied in their readiness to remate following a first mating with the tester male: California females remated more readily while Greece and Taiwan females mated less readily than average (Figure 5, Table S5). Male genotype also affected remating: California males were more consistently successful in these second matings, while Greece males were less successful on average (Figure 5, Table S5). In contrast, we detected no effect of male perfuming status on second-male premating success (Figure 5, Table S5).

**Figure 5.**
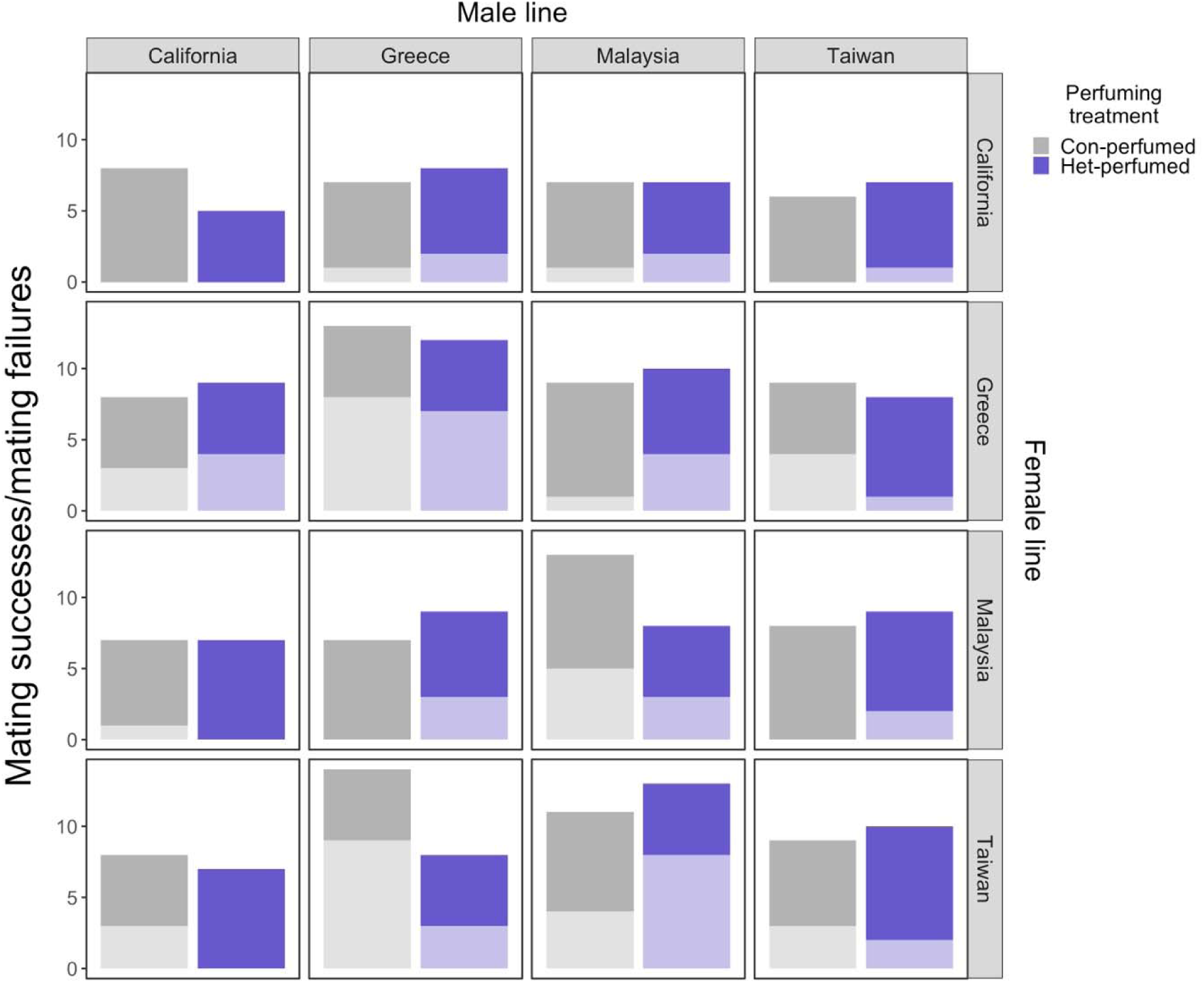
Counts of mating successes (dark shading) and mating failures (light shading) differs across male (column) and female (row) genotypes (Table S5). Results are shown separately for conspecific-perfumed males (grey) and heterospecific-perfumed males (purple) within each male x female panel. **Alt text:** A matrix of graphs representing mating successes vs. failures across all possible male by female combinations, divided according to conspecific- or heterospecific-perfumed treatments. Data are represented with stacked bars, where the lower bar represents the count of mating failures and the upper bar represents mating successes.

### 3.5 Female genotypes vary in their overall use of sperm from the second male, and in whether they adjust sperm use according to second-male perfuming status

We assessed how paternity of second-males (P2) was affected by genotype and by our perfuming manipulation. We found strong and significant differences in second-male paternity depending on both female and second-male genotype, female by male genotype interaction effects, and female genotype by male genotype by treatment interaction effects. In particular, Taiwan males gained higher P2 than average, while Greece and Malaysia males had reduced paternity share (Figure 6, Table 1). In comparison, Greece females exhibited a higher P2 than average, while Taiwan females exhibited a lower P2 than average, across all genotype pairs (Figure 6, Table 1). Despite these general effects, second-male paternity share also varied substantially depending on the particular male-female combination. For instance, Greece females exhibit higher average P2 when paired with Taiwan second males, while Taiwan females reduce P2 when paired with Malaysia second males (Figure 6, Table 1).

**Figure 6.**
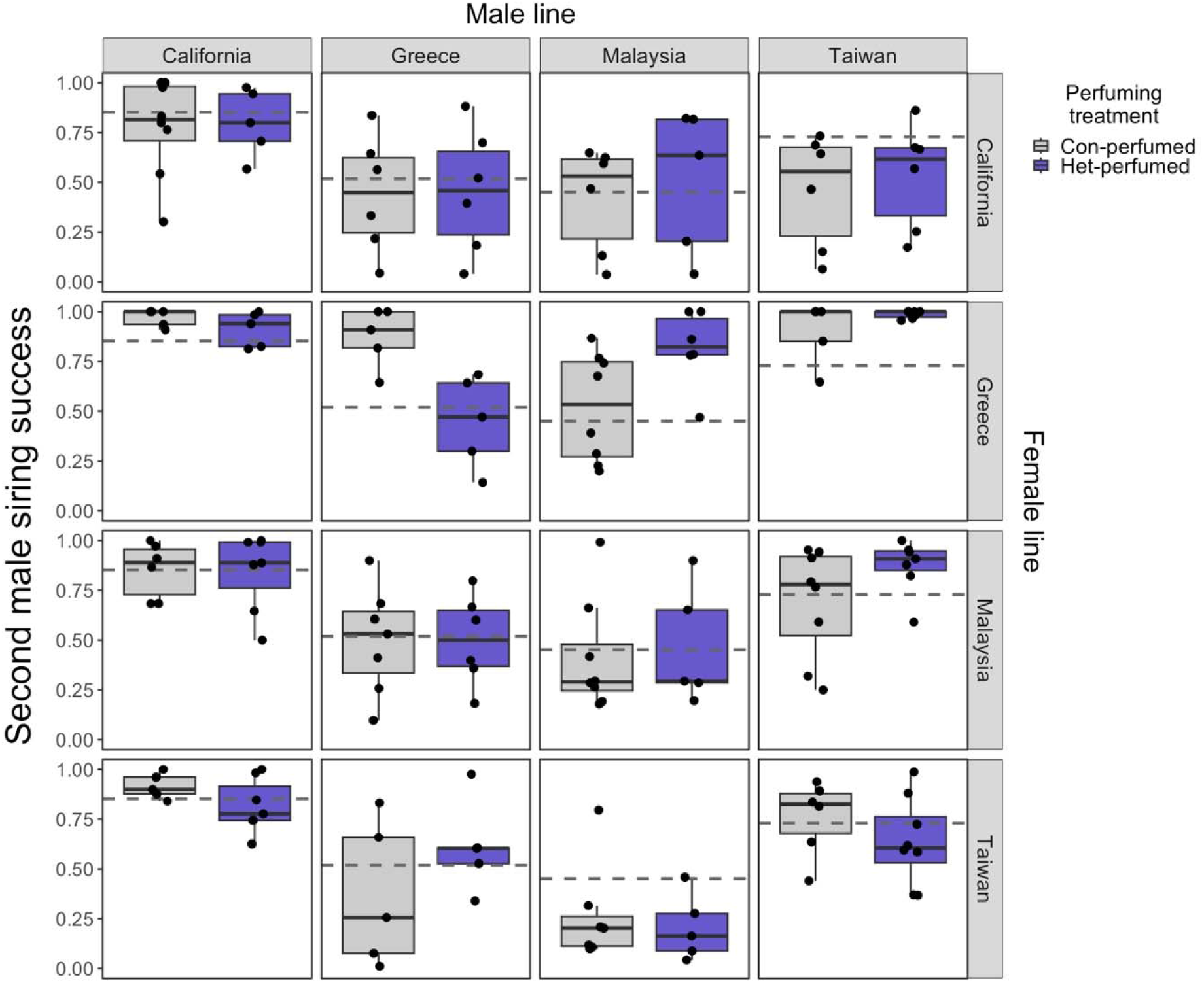
Female use of second male sperm (P2) varies across female genotypes, male genotypes, and female by male genotype combinations (Table 1). P2 also varies depending on male perfuming status, as shown in grey (conspecific-perfumed males) and purple (heterospecific-perfumed males). The dashed line indicates the average P2 for each male line. Box plots show the median (central line) and first and third quartiles; whiskers extend to the smallest and largest values (not including outliers >1.5x the inter-quartile range). **Alt text:** A matrix of graphs representing second male siring success across all possible male by female combinations, divided according to conspecific- or heterospecific-perfumed treatments. Data are represented using box plots and individual data points.

**Table 1.**
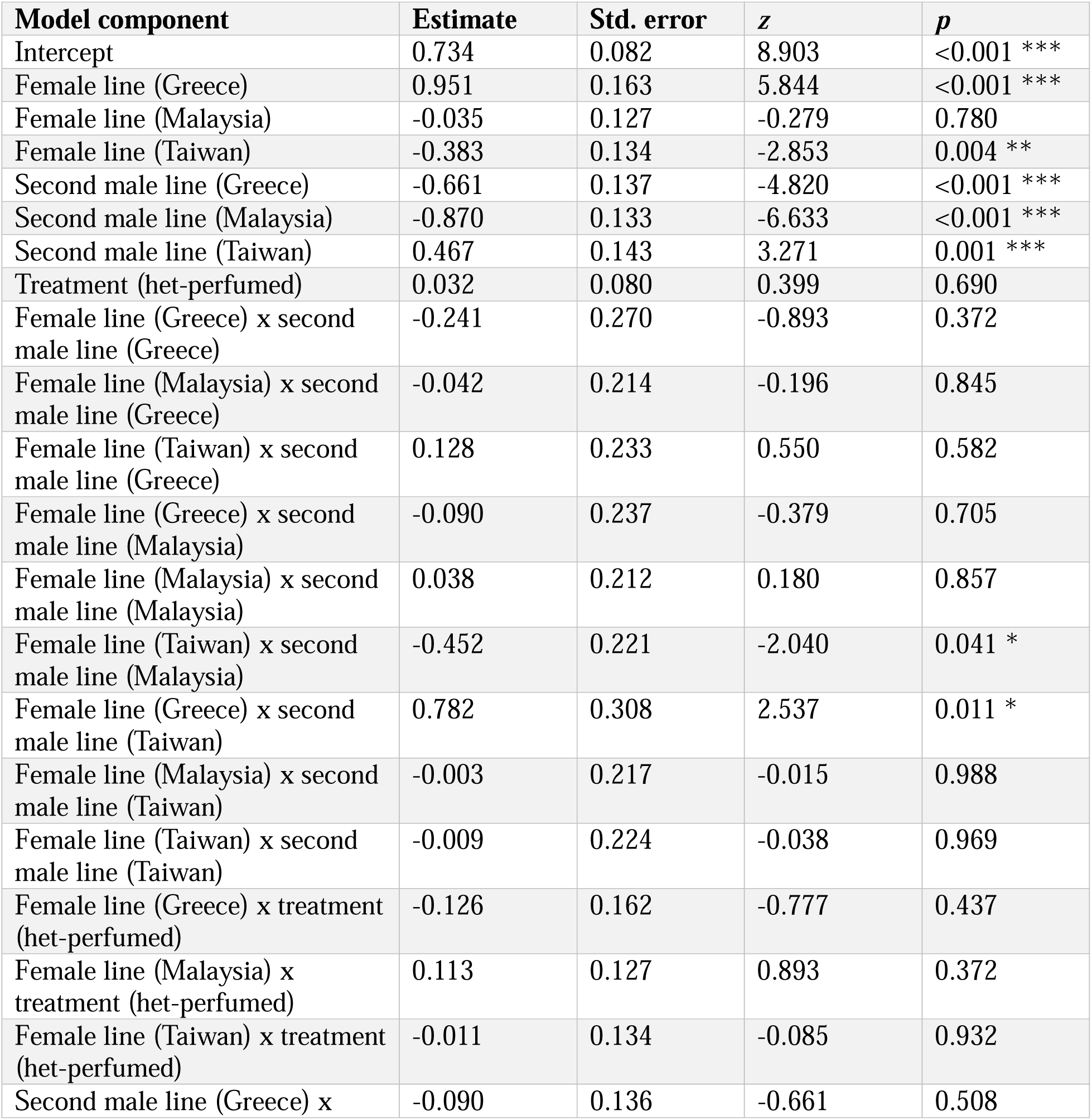
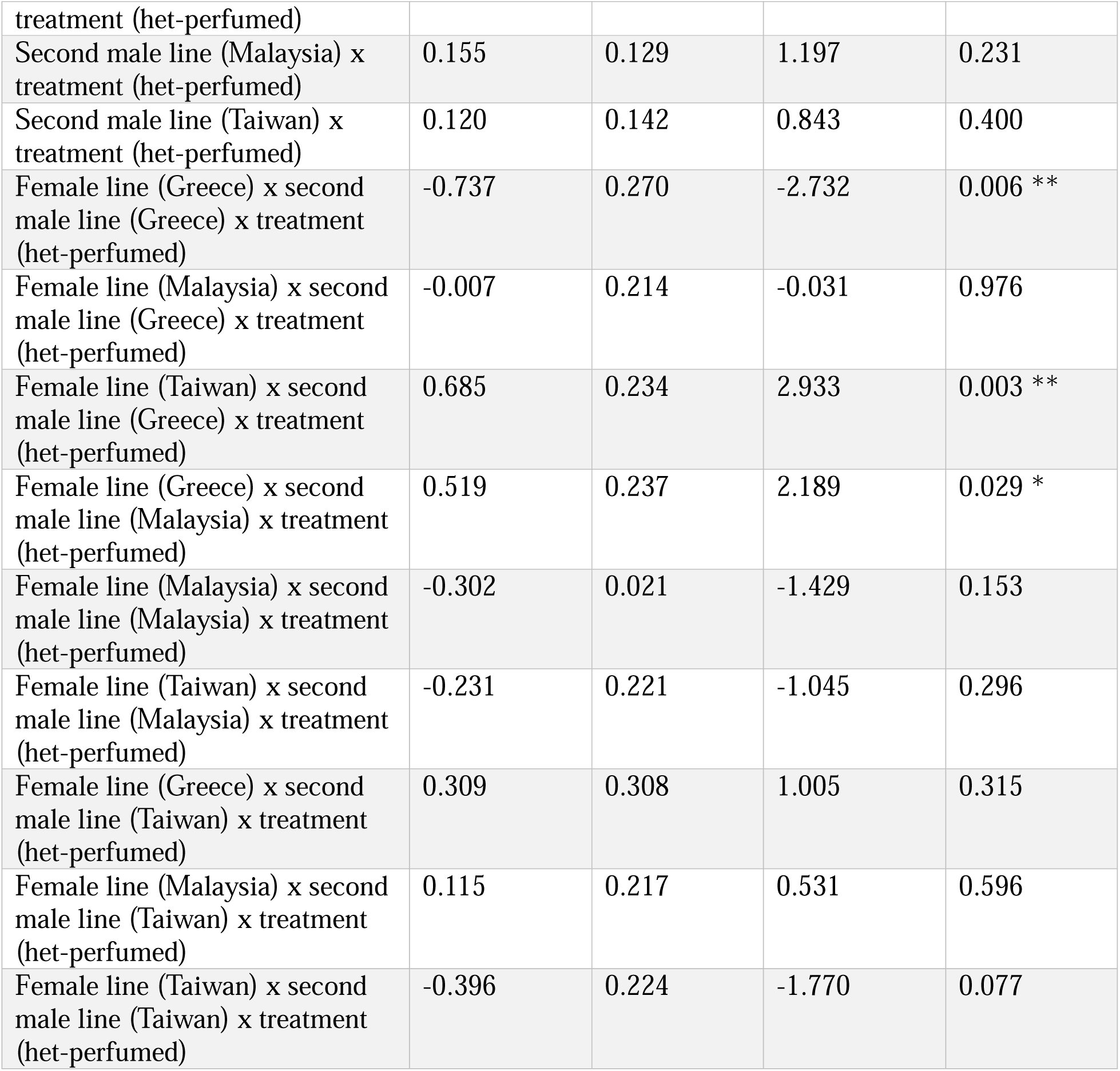
Specific genotype, treatment, and interaction effects on second male paternity share (P2). Shown is the beta-binomial generalized linear model of variation in second male paternity share (P2), depending on female line, second male line, treatment group, and all possible interactions. California females, California males, and conspecific-perfumed males are incorporated into the model as a baseline. **Alt text:** A table containing the output of our beta-binomial generalized linear model. Our model includes significant terms for at least some components of female genotype, second male genotype, female genotype by second male genotype, and female genotype by second male genotype by perfuming treatment.

We also found that females adjust sperm use according to male genotype and male perfuming treatment, including in female genotype-specific ways. In particular, female genotypes vary in whether and how sperm use patterns were affected by perfuming. For instance, Greece females use less sperm from Greece males specifically when those males are heterospecific-perfumed (Figure 6, Table 1). In contrast, Taiwan females use more sperm from heterospecific-perfumed Greece males (Figure 6, Table 1). Greece females also use more sperm from Malaysia males when they are perfumed by heterospecifics (Figure 6, Table 1). In comparison, California and Malaysia females do not appear to respond to male CHC perfuming when making sperm use choices.

### 3.6 Both pre- and postmating choices can be affected by premating behavior, but sperm length shows the most consistent association with patterns of preferential sperm use

We next assessed correlations between datasets to better understand how these variables might influence one another. First, we evaluated the relationship between second-male courtship effort and second-male mating success—i.e., is premating courtship intensity associated with premating success? We did not see a consistent association for either conspecific-perfumed males (Figure S4a; Spearman’s rank correlation ρ = 0.308, *p* = 0.246) or heterospecific-perfumed males (Figure S4b; Spearman’s rank correlation ρ = 0.174, *p* = 0.519). Instead, readiness to remate seems to be more clearly associated with individual female line differences. For instance, California females tend to remate most readily, and Greece and Taiwan least readily, regardless of the courtship intensity of the male line they are paired with (Figure S4a,b).

Second, we assessed the relationship between second male courtship effort and paternity share (P2)—i.e., do females use premating signals to make postmating decisions? We found that when males are perfumed by conspecifics, male courtship effort is significantly correlated with siring success (Figure 7a; Spearman’s rank correlation ρ = 0.535, *p* = 0.035). When males are perfumed by heterospecifics, the correlation is positive but not significant (Figure 7b; Spearman’s rank correlation ρ = 0.353, *p* = 0.180). This weak positive trend between courtship effort and P2 appears to be largely driven by California males, which are the most active courters and also received the greatest share of paternity (Figure 7a,b), an association which is unclear in the other male lines. Thus, we infer that the relationship between these variables is likely influenced by additional factors not measured here; for instance, Greece males, which have low siring success, have been shown to exhibit especially long interpulse intervals during courtship song, which are less preferred by *D. melanogaster* females (Kyriacou & Hall 1982, Pischedda et al. 2014a).

**Figure 7.**
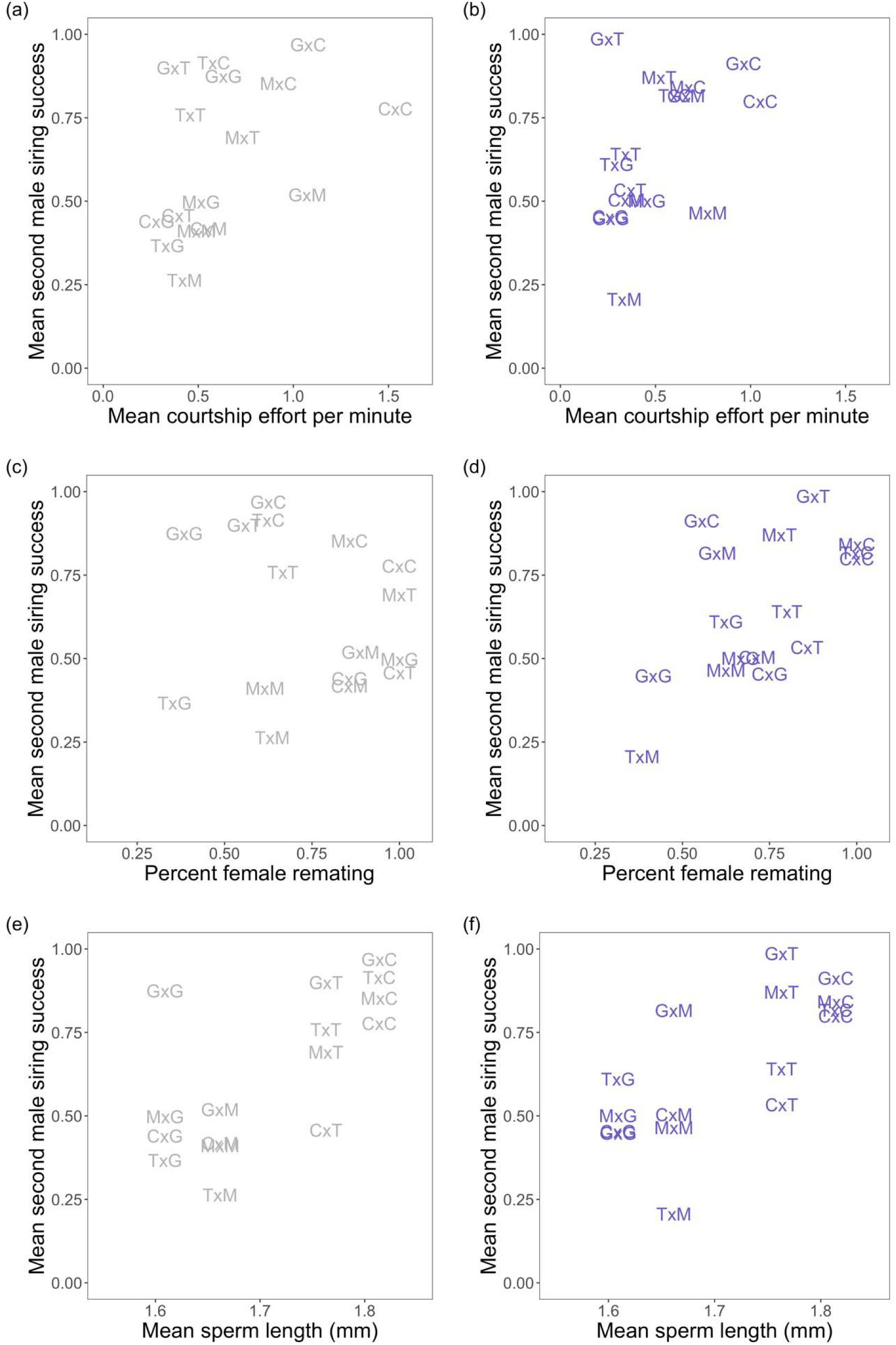
Second male courtship effort is significantly correlated with siring success when males are perfumed by conspecifics (a),but is not significant when perfumed by heterospecifics (b). Female remating rate is not correlated with second male siring success when males are conspecific-perfumed (c) but is correlated when males are heterospecific-perfumed (d). Sperm length is significantly correlated with siring success for both conspecific-perfumed (e) and heterospecific-perfumed males (f). Crosses are labeled as female x male genotype (G, Greece; T, Taiwan; C, California; and M, Malaysia). **Alt text:** Scatterplots representing the correlation between datasets. Points are represented using the first letter of each genotype in that cross (for example, C x G is California female by Greece male). Data is divided between conspecific-perfumed and heterospecific-perfumed males. Sperm length and second male siring success show the clearest pattern of correlation.

We next assessed the relationship between female readiness to remate and second-male paternity share—i.e., is premating choosiness associated with postmating choosiness? We found no correlation among these factors for conspecific-perfumed males (Figure 7c; Spearman’s rank correlation ρ = −0.095, *p* = 0.726), and a slight association for experiments with heterospecific-perfumed males (Figure 7d; Spearman’s rank correlation ρ = 0.502, *p* = 0.048). These weak or absent associations are consistent with heterogenous effects we observe among female lines. For instance, while Greece females showed relatively high premating choosiness (remating in only 59% of trials) and the highest average P2 of any genotype (consistent with strong second male preference), Taiwan females exhibit a similar premating choosiness (60% remating frequency) but significantly lower average P2 (indicating no consistent postmating preference for first or second male sperm) (Figure 7c,d). This relationship therefore appears to vary substantially depending upon which of the four genotypes is considered.

Finally, we examined the relationship between second-male paternity share and reproductive trait variation among lines. Of RT mass (both male and female), SR length, testis length, sperm length, and SR length minus sperm length, only sperm length and MRT mass were significantly associated with P2. Of these two, MRT mass was significantly negatively correlated with P2 among heterospecific-perfumed males (Figure S4d; Spearman’s rank correlation ρ = −0.679, *p* = 0.004), but not among conspecific-perfumed males (Figure S4c; Spearman’s rank correlation ρ = −0.449, *p* = 0.081). These relationships appear to be substantially influenced by Taiwan males, which have high P2 but comparatively low RT mass. In comparison, the relationship between sperm length and P2 was consistently and strongly positive for both conspecific-perfumed (Figure 7e; Spearman’s rank correlation ρ = 0.631, *p* = 0.009) and heterospecific-perfumed groups (Figure 7f; Spearman’s rank correlation ρ = 0.728, *p* = 0.001). Second males with longer sperm experience greater siring success. Specifically, when the second male’s sperm is longer than the tester male (as for California or Taiwan second males), P2 is generally higher; whereas when the second male sperm is shorter than the tester male (as for Malaysia and Greece), P2 is generally lower. Together, these data indicate that females preferentially use longer sperm—a generic preference that appears as last male precedence when the second male’s sperm is longer than the first (tester) male’s sperm, but can override last male precedence when the second male’s sperm is relatively shorter.

## 4 Discussion

In this study, we sought to understand whether and how females make plastic choices about sperm use. We found that sperm use patterns vary based on male genotype, female genotype, their interaction, as well as based on male pheromone profile (even when male identity is the same). Among these effects, male sperm length, specifically the relative difference in sperm length between the first and second male, appears to play a particularly important role in determining female sperm use patterns. While sperm length has been recognized as an important component of male-male competition (e.g., Immler et al. 2011, Lüpold & Pitnick 2018, Snook 2005), our analysis enables us to infer that the preference for longer sperm is also female-mediated, context-dependent, and controlled by factors other than the size of the female sperm storage organs, as we expand upon below.

### 4.1 Female sperm use choices vary among lines in their direction and magnitude, and in the criteria used to evaluate males

Consistent with previous work in *D. melanogaster*, we observe clear evidence for sperm choice mediated by females (Chow et al. 2010, Clark & Begun 1998, Clark et al. 1999, Miller & Pitnick 2002). Female lines show patterns of sperm use in control crosses (with their own unmanipulated second males) that significantly shift in response to changing either second-male genotype or perfuming treatment or both (depending on the female line). In addition, both first and second males have the potential to shape some paternity outcomes via plastic allocation during the transfer of sperm, including adjusting ejaculate composition based on the identity of the female. In some cases, it is difficult to differentiate between these male or female plastic effects; for instance, our male by female genotype interaction effects on paternity share could be shaped by both cryptic male and cryptic female choice. However, our design specifically aimed to include effects where the influence of male plasticity should be minimized. In particular, to attribute perfuming effects on paternity to cryptic male choice, males would have to plastically adjust their ejaculate based on one minute of exposure to conspecific vs. heterospecific males, but only when mating with certain female genotypes. A more parsimonious and biologically plausible inference is that females are adjusting paternity—exercising choice—in response to male pheromone differences. Thus, in addition to the effects of male reproductive plasticity, our results underscore the potential for cryptic female choice to influence genotypic representation among the resulting offspring, and to generate sexual selection on males (Pitnick et al. 2001).

We found substantial and consistent variation among our four female lines in the direction and magnitude of preferential sperm use patterns. *D. melanogaster* is conventionally described as having second (or last) male sperm precedence (Lefevre & Jonsson 1962, Pischedda & Rice 2012). However, depending on the specific male-female combination, our estimated preference for the sperm of second-males ranged from 0.2 to 0.95. Values below 0.5 (e.g., Taiwan females with Malaysia males) indicate consistent first-male precedence in these specific male-female combinations. Moreover, female lines differ in how strongly they differentiate among sperm donors. For example, with the exception of California males (which they strongly prefer; average P2 = 0.79), California females do not consistently differentiate between the first and second males’ sperm when paired with the other three second male lines in our experiment (i.e., mean P2 is approximately 0.5, with a broad range of values among individual CA females; Figure 6, Table S6).

We also clearly observed among-line variation in the characteristics that females use to evaluate males as reproductive donors. In particular, Greece and Taiwan females modulate their use of second male sperm in response to perfuming status, albeit in complex ways. For example, Greece females decrease their use of Greece sperm when these males are heterospecific-perfumed, but Taiwan females increase their use of Greece sperm after the same treatment (Figure 6, Table S7). In comparison, male perfuming appears to have no influence on cryptic choice by California or Malaysia females. This diversity in sperm choice patterns depending on intrinsic (female genotype) and extrinsic (perfumed status) factors points to a much greater heterogeneity of sperm use decision-making among *D. melanogaster* females than simple last male precedence (see also Laturney et al. 2018). Given the broad geographic origin of these lines, differences in the magnitude and direction of female choices could reflect reproductive or other differences among lines that operated prior to their collection and lab establishment.

### 4.2 Cryptic female choice is strongly shaped by male traits, especially sperm length

Within the diversity of individual sperm use criteria, however, we did observe several broad patterns. In particular, every female line differentiated among male genotypes in at least one postmating experimental context, indicating that one or more aspects of male genotype are important in making paternity allocation decisions. Our analysis also identifies specific male traits that likely mediate these male genotype effects. As noted above, some female lines adjust sperm use in response to perfuming treatment, indicating that male cuticular hydrocarbon profile affects their postmating choices. Among conspecific-perfumed second males, paternity share is also associated with courtship effort, suggesting that females may prefer the sperm of male genotypes who court more, although this effect is shaped largely by California males. Most strikingly, however—and more consistently than either chemosensory or courtship traits—we find that sperm length is positively associated with preferential sperm use across many contexts and conditions. The majority of female lines preferentially used the second-male’s sperm when that sperm was longer than the first-male’s sperm (i.e., when California or Taiwan was the second male). This result is consistent with previous studies demonstrating the increased competitive ability of long sperm (e.g., Miller & Pitnick 2002). However, if this process were solely mediated by sperm length, we would expect paternity outcomes to be consistent across female genotypes. Instead, we see significant variation in male siring success among female lines and perfuming treatments; for instance, Taiwan males show increased siring success when mating with Greece females compared to California females (Figure 6, Table 1), indicating that females also play a role in shaping these outcomes. Moreover, in several cases (for example, when Taiwan females mate with Malaysia males—see Figure 6) female lines subverted the expected precedence for last males’ sperm specifically when the first (GFP tester) male had longer sperm than the second male (i.e. Greece and Malaysia males). That is, our results indicate that ‘long-sperm precedence’, in addition to ‘last-male sperm precedence’ (Lefevre & Jonsson 1962), jointly contribute to female sperm use patterns in *D. melanogaster*.

Finally, although our data indicate diverse criteria for female sperm choice overlaid on a generic preference for longer sperm, it also indicates that these patterns are not explained by at least one studied mechanism of female sperm choice—SR length (Miller & Pitnick 2002). SR length has been proposed as a primary mechanism via which females discriminate between sperm of different lengths (Pitnick et al. 1999); in particular, females with longer SRs are thought to be most selective because they can better discriminate between the sperm length of the first and second male. However, while we observe clear variation in female discrimination of sperm based on length—e.g., Greece females exhibit generally high P2, regardless of length, whereas California females only exhibit high P2 for their own (long-spermed) males (Figure 6, Table 1)— we do not observe significant variation in SR length among our female lines (Figure 3, Table S3, Table S2). Thus, SR length is not the mechanism by which these diverse *D. melanogaster* female lines discriminate against short sperm in our experiments.

### 4.3 Evolutionary implications of the observed variation in female sperm choice

Consistent patterns in choice criteria among female genotypes provide an opportunity to evaluate the nature of past drivers of female choice in our system, including hypotheses where cryptic female choice is not beneficial to females versus where it has specific fitness benefits. For example, one proposal is that female sperm use patterns result primarily from manipulation by males; that is, cryptic female choice is the product of sexual conflict rather than providing an adaptive benefit to females. In this case, female sperm use patterns are expected to be random with respect to actual measures of male quality. Our data do not convincingly support this hypothesis, given the general postmating female preferences we observe for males who have the longest sperm and, under some conditions, for males that court frequently.

Alternatively, Birkhead (1998) proposed that female sperm choice might function to bias sperm use against the sperm of closely related males. However, our findings are not consistent with inbreeding avoidance as a primary function of cryptic female choice. If that were the case, second-male success (against the unrelated tester first male) should consistently be lower in crosses with males from the same line as females, compared to males from different lines. However, female lines differed in their choice of own male sperm, ranging from consistent high preference to low preference.

Apart from inbreeding avoidance, other hypothesized functions of sperm choice include female benefits from sire discrimination (e.g., choice of high-quality males [‘good genes’ hypothesis; Eberhard 1996], or choice of attractive males [‘sexy sons’ hypothesis; Weatherhead & Robertson 1979]). We find that females have a generic preference for longer sperm (Figure 7e,f); in this case, choosing longer-spermed males increases the frequency of longer-spermed (i.e. ‘sexier’) sons, which themselves could benefit from the same generic preference. As studies have shown that long sperm have a competitive advantage in displacing shorter sperm (Lüpold et al. 2012, Lüpold et al. 2016), this preference likely also aligns specifically with higher reproductive fitness in males. Thus, our data support the hypothesis that females could gain indirect fitness benefits from exercising cryptic choice based on specific male traits like sperm length.

The differences we observe among female lines in the criteria they used for sperm choice also have implications for the expected future drivers of postmating reproductive evolution. Some of our lines altered sperm use patterns based on contact pheromones (Greece and Taiwan), indicating that pheromone bouquet can be a criterion for evaluating preferred sires. Other lines (California and Malaysia) were insensitive to pheromone manipulation in their sperm use choices. Variation in choice criteria makes it more challenging for males to tailor specific traits to satisfy, or manipulate (Ryan 1990), a broad range of female biases. Moreover, when the specific preferences of females differ according to male trait values and male trait identity, there is no single most preferred male genotype, and variation should be maintained at male traits. This expectation is consistent with our observation, and with previous analyses (e.g., Clark et al. 1999, Chow et al. 2010, Miller & Pitnick 2002), showing that the outcomes of cryptic female choice are demonstrably influenced by male by female interactions.

That these criteria for female sperm choice can also vary behaviorally is particularly evident in the case of Greece females. When sequentially mated with males from other lines, Greece females uniformly favor whichever genotype (tester versus second male) has the longer sperm. However, when mated with their own conspecific-perfumed second males, Greece females preferentially use this sperm (Figure 6), even though it is significantly shorter than the first (tester) male (Figure 3). In this case, the female preference for local males appears to override their generic preference for longer sperm. We know that this decision is both female-mediated and based on chemosensory assessment, because it is reversed when Greece females are paired with heterospecific-perfumed Greece males. Greece females instead favor the longer (tester male) sperm in these pairings (Figure 6), even though these Greece males are identical in every sense to conspecific-perfumed males—except for their CHC profile. Flexible changes in preference, such as this case where females specifically prefer own males despite generic preferences for longer sperm, is another potential mechanism by which sexually selected variation could be maintained in males.

## 5 Conclusion

The direction and magnitude of female-mediated sperm choice, as well as the male traits influencing these choices, varies significantly among four female lines of *D. melanogaster*. This diversity in female sperm choices is overlaid on a generic preference for longer sperm, suggesting that paternity decisions may ultimately benefit female fitness indirectly through the production of higher-quality offspring or more attractive offspring. Interestingly, at least one proposed proximate mechanism of female sperm choice—SR length—does not account for variation in observed sperm use. Diversity in cryptic female choices—both in the specific modalities used to assess and choose among males, and in the plasticity of these choices—may explain how variation in male postmating sexually selected traits is maintained over time.

## Data availability statement

The data and code underlying this article are available in the Dryad Digital Repository, at https://dx.doi.org/[doi].

## Author contributions

BP and LCM conceived and designed the experiment. BP performed the experiment and statistical analyses. JYY performed the chemical analysis. BP and LCM wrote the paper. All authors read and provided feedback on the manuscript.

## Acknowledgments

We thank Stuart J. Macdonald for providing the DSPR founder lines used in this study, and Brandon S. Cooper for providing the *Drosophila yakuba* line. We thank Sung Nichin for assistance scoring courtship behaviors. We thank Alison Pischedda for advice on the perfuming protocol. All GCMS measurements were performed in the Univ. of Hawai i at Mānoa Microbial Genomics and Analytical Laboratory core. We thank Matthew W. Hahn, Kimberly A. Rosvall, and Michael J. Wade for their helpful comments on the manuscript. The research was supported by Indiana University Department of Biology funding to LCM and BP, as well as the NIH Common Themes in Reproductive Diversity (CTRD) training grant funding to BP.

