## Supplementary Material for "Long-sperm precedence and other cryptic female choices in *Drosophila melanogaster*"

| Table S1. Variation in CHC composition across male genotypes and perfuming treatments, assessed using a multivariate analysis of variance (MANOVA) on the first principal components from a PCA of 16 compounds generated from GCMS (Table S5).  Alt text: A table containing the output of our MANOVA on the first four principal components from a PCA. The first three PCs show significant effects of second male genotype, and PC3 and PC4 show significant effects of perfuming treatment. | | | | | |
| --- | --- | --- | --- | --- | --- |
| PC | **Model component** | **Sum sq.** | **Mean sq.** | **F** | ***p*** |
| PC1 | Genotype | 274.738 | 91.579 | 334.623 | <0.001 *** |
|  | Treatment | 0.015 | 0.015 | 0.053 | 0.819 |
|  | Residuals | 9.579 | 0.274 |  |  |
| PC2 | Genotype | 226.290 | 75.430 | 93.999 | <0.001 *** |
|  | Treatment | 0.144 | 0.144 | 0.180 | 0.674 |
|  | Residuals | 28.086 | 0.802 |  |  |
| PC3 | Genotype | 27.519 | 9.173 | 42.797 | <0.001 *** |
|  | Treatment | 1.994 | 1.994 | 9.304 | 0.004 ** |
|  | Residuals | 7.502 | 0.214 |  |  |
| PC4 | Genotype | 1.659 | 0.553 | 2.730 | 0.0585 |
|  | Treatment | 8.346 | 8.346 | 41.201 | <0.001 *** |
|  | Residuals | 7.090 | 0.203 |  |  |

| Table S2. CHC means and standard deviations for conspecific-perfumed (C) and heterospecific-perfumed (H) males.  Alt text: A table containing the means and standard deviations of CHC compounds across male genotype and perfuming treatment. | | | | | | | | |
| --- | --- | --- | --- | --- | --- | --- | --- | --- |
| CHC | **Line** | | | | | | | |
|  | **California** | | **Greece** | | **Malaysia** | | **Taiwan** | |
|  | **C** | **H** | **C** | **H** | **C** | **H** | **C** | **H** |
| nC21(11.08) | 0.164  ± 0.014 | 0.177  ± 0.010 | 0.170  ± 0.016 | 0.198  ± 0.025 | 0.265  ± 0.021 | 0.256  ± 0.010 | 0.254  ± 0.016 | 0.260  ± 0.025 |
| C22:1(11.4) | 0.038  ± 0.003 | 0.045  ± 0.005 | 0.045  ± 0.004 | 0.056  ± 0.009 | 0.072  ± 0.006 | 0.072  ± 0.004 | 0.099  ± 0.005 | 0.094  ± 0.014 |
| cVA(11.47) | 0.161  ± 0.045 | 0.402  ± 0.227 | 0.063  ± 0.007 | 0.103  ± 0.070 | 0.609  ± 0.107 | 0.689  ± 0.121 | 1.433  ± 0.230 | 1.575  ± 0.175 |
| nC22(11.51) | 0.148  ± 0.009 | 0.184  ± 0.017 | 0.109  ± 0.009 | 0.135  ± 0.017 | 0.175  ± 0.012 | 0.194  ± 0.008 | 0.189  ± 0.013 | 0.196  ± 0.012 |
| 9-T(11.84) | 0.114  ± 0.038 | 0.123  ± 0.037 | 0.403  ± 0.013 | 0.373  ± 0.117 | 0.391  ± 0.057 | 0.346  ± 0.037 | 0.398  ± 0.089 | 0.429  ± 0.085 |
| 7-T(11.87) | 2.406  ± 0.197 | 2.713  ± 0.154 | 3.872  ± 0.256 | 4.330  ± 0.459 | 4.574  ± 0.340 | 4.294  ± 0.288 | 6.031  ± 0.346 | 6.094  ± 0.629 |
| 5-T(11.92) | 0.174  ± 0.017 | 0.160  ± 0.022 | 0.296  ± 0.013 | 0.287  ± 0.033 | 0.354  ± 0.024 | 0.294  ± 0.023 | 0.394  ± 0.027 | 0.371  ± 0.040 |
| nC23(11.94) | 0.826  ± 0.065 | 0.818  ± 0.050 | 1.172  ± 0.084 | 1.292  ± 0.162 | 1.000  ± 0.065 | 0.907  ± 0.043 | 1.053  ± 0.077 | 1.077  ± 0.098 |
| MeC24(12.75) | 0.228  ± 0.025 | 0.212  ± 0.021 | 0.095  ± 0.012 | 0.089  ± 0.011 | 0.255  ± 0.011 | 0.210  ± 0.025 | 0.324  ± 0.024 | 0.293  ± 0.023 |
| 9-P(12.83) | 0.057  ± 0.008 | 0.064  ± 0.003 | 0.149  ± 0.008 | 0.153  ± 0.021 | 0.085  ± 0.007 | 0.077  ± 0.011 | 0.151  ± 0.007 | 0.144  ± 0.021 |
| 7-P(12.87) | 0.367  ± 0.041 | 0.379  ± 0.023 | 1.649  ± 0.094 | 1.591  ± 0.189 | 0.345  ± 0.025 | 0.298  ± 0.035 | 0.538  ± 0.034 | 0.503  ± 0.078 |
| 5-P(12.92) | 0.000  ± 0.000 | 0.000  ± 0.000 | 0.096  ± 0.007 | 0.087  ± 0.009 | 0.000  ± 0.000 | 0.000  ± 0.000 | 0.000  ± 0.000 | 0.000  ± 0.000 |
| MeC26(14.03) | 0.548  ± 0.047 | 0.614  ± 0.041 | 0.709  ± 0.034 | 0.771  ± 0.107 | 0.476  ± 0.019 | 0.477  ± 0.038 | 0.515  ± 0.022 | 0.650  ± 0.046 |
| 7-H(14.21) | 0.000  ± 0.000 | 0.019  ± 0.007 | 0.150  ± 0.014 | 0.176  ± 0.025 | 0.000  ± 0.000 | 0.000  ± 0.000 | 0.000  ± 0.000 | 0.005  ± 0.001 |
| nC27(14.32) | 0.037  ± 0.011 | 0.048  ± 0.010 | 0.058  ± 0.016 | 0.080  ± 0.016 | 0.030  ± 0.003 | 0.028  ± 0.003 | 0.069  ± 0.011 | 0.056  ± 0.010 |
| MeC28(15.9) | 0.201  ± 0.022 | 0.203  ± 0.025 | 0.272  ± 0.045 | 0.280  ± 0.045 | 0.333  ± 0.012 | 0.250  ± 0.044 | 0.392  ± 0.024 | 0.395  ± 0.041 |

| Table S3. Variation in female and male postmating reproductive traits, as assessed in linear models. GFP-labelled females and GFP-labelled males are incorporated into the models as a baseline.  Alt text: A table containing the output of our linear models. We see significant variation in morphology across genotypes for most traits, with notable exceptions being seminal receptacle length and spermtheca area. | | | | | |
| --- | --- | --- | --- | --- | --- |
| Response | **Model component** | **Estimate** | **Std. error** | ***t*** | ***p*** |
| Male thorax length | Intercept | 0.795 | 0.008 | 105.693 | <0.001 *** |
|  | Male line (California) | -0.006 | 0.015 | -0.425 | 0.675 |
|  | Male line (Greece) | 0.042 | 0.015 | 2.790 | 0.011 * |
|  | Male line (Malaysia) | -0.031 | 0.015 | -2.060 | 0.053 |
|  | Male line (Taiwan) | -0.076 | 0.015 | -5.063 | <0.001 *** |
| Testis length | Intercept | 2.768 | 0.726 | 3.813 | 0.001 ** |
|  | Male thorax length | -0.174 | 0.912 | -0.191 | 0.851 |
|  | Male line (California) | 0.227 | 0.062 | 3.690 | 0.002 ** |
|  | Male line (Greece) | 0.246 | 0.072 | 3.402 | 0.003 ** |
|  | Male line (Malaysia) | -0.293 | 0.068 | -4.333 | <0.001 *** |
|  | Male line (Taiwan) | -0.299 | 0.093 | -3.221 | 0.004 ** |
| Sperm length | Intercept | 1.729 | 0.069 | 25.237 | <0.001 *** |
|  | Male dry body mass (without MRT) | -0.072 | 0.289 | -0.250 | 0.805 |
|  | Male line (California) | 0.101 | 0.014 | 7.025 | <0.001 *** |
|  | Male line (Greece) | -0.105 | 0.017 | -6.236 | <0.001 *** |
|  | Male line (Malaysia) | -0.049 | 0.015 | -3.330 | 0.004 ** |
|  | Male line (Taiwan) | 0.053 | 0.017 | 3.060 | 0.006 ** |
| Male dry body mass (without MRT) | Intercept | 0.236 | 0.006 | 42.550 | <0.001 *** |
|  | Male line (California) | -0.006 | 0.011 | -0.564 | 0.579 |
|  | Male line (Greece) | -0.308 | 0.011 | -2.783 | 0.011 * |
|  | Male line (Malaysia) | 0013 | 0.011 | 1.217 | 0.238 |
|  | Male line (Taiwan) | 0.034 | 0.011 | 3.094 | 0.006 ** |
| MRT dry mass | Intercept | 0.019 | 0.006 | 3.034 | 0.007 ** |
|  | Male dry body mass (without MRT) | 0.031 | 0.026 | 1.165 | 0.258 |
|  | Male line (California) | -0.001 | 0.001 | -0.736 | 0.471 |
|  | Male line (Greece) | 0.004 | 0.002 | 2.619 | 0.017 * |
|  | Male line (Malaysia) | 0.002 | 0.001 | 1.113 | 0.280 |
|  | Male line (Taiwan) | -0.005 | 0.002 | -2.994 | 0.007 ** |
| Female thorax length | Intercept | 0.884 | 0.008 | 112.185 | <0.001 *** |
|  | Female line (California) | -0.002 | 0.016 | -0.102 | 0.920 |
|  | Female line (Greece) | 0.022 | 0.016 | 1.409 | 0.174 |
|  | Female line (Malaysia) | -0.048 | 0.016 | -3.072 | 0.006 ** |
|  | Female line (Taiwan) | -0.039 | 0.016 | -2.488 | 0.022 * |
| SR length | Intercept | 1.827 | 1.210 | 1.510 | 0.147 |
|  | Female thorax length | 0.317 | 1.368 | 0.232 | 0.819 |
|  | Female line (California) | 0.027 | 0.096 | 0.284 | 0.780 |
|  | Female line (Greece) | -0.055 | 0.101 | -0.547 | 0.591 |
|  | Female line (Malaysia) | -0.192 | 0.117 | -1.639 | 0.118 |
|  | Female line (Taiwan) | -0.034 | 0.110 | -0.309 | 0.760 |
| ST area | Intercept | 0.002 | 0.003 | 0.753 | 0.461 |
|  | Female thorax length | 0.003 | 0.004 | 0.717 | 0.482 |
|  | Female line (California) | 0.000 | 0.000 | -0.515 | 0.613 |
|  | Female line (Greece) | 0.000 | 0.000 | 1.592 | 0.128 |
|  | Female line (Malaysia) | 0.000 | 0.000 | -1.168 | 0.257 |
|  | Female line (Taiwan) | 0.000 | 0.000 | 1.156 | 0.262 |
| Female dry body mass (without FRT) | Intercept | 0.298 | 0.005 | 56.450 | <0.001 *** |
|  | Female line (California) | -0.010 | 0.011 | -0.913 | 0.372 |
|  | Female line (Greece) | -0.021 | 0.011 | -2.018 | 0.057 |
|  | Female line (Malaysia) | -0.024 | 0.011 | -2.260 | 0.0351 * |
|  | Female line (Taiwan) | 0.059 | 0.011 | 5.579 | <0.001 *** |
| FRT dry mass | Intercept | -0.067 | 0.052 | -1.278 | 0.217 |
|  | Female dry body mass (without FRT) | 0.481 | 0.175 | 2.755 | 0.013 * |
|  | Female line (California) | 0.023 | 0.008 | 2.689 | 0.015 * |
|  | Female line (Greece) | -0.004 | 0.009 | -0.405 | 0.690 |
|  | Female line (Malaysia) | 0.017 | 0.009 | 1.860 | 0.078 |
|  | Female line (Taiwan) | -0.051 | 0.013 | -3.844 | 0.001 ** |

| Table S4. Variation in second male courtship effort (courtship behaviors per minute) depending on female line, second male line, treatment group, and the interaction between female line and second male line, as assessed in a linear model. California females, California males, and conspecific-perfumed males are incorporated into the model as a baseline.  Alt text: A table containing the output of our linear model. Our model includes significant terms for at least some components of female genotype and second male genotype, but no significant effects from perfuming treatment or the interaction between female genotype and male genotype. | | | | |
| --- | --- | --- | --- | --- |
| Model component | **Estimate** | **Std. error** | ***t*** | ***p*** |
| Intercept | 0.579 | 0.037 | 15.837 | <0.001 *** |
| Female line (Greece) | 0.082 | 0.063 | 1.304 | 0.194 |
| Female line (Malaysia) | 0.055 | 0.064 | 0.866 | 0.388 |
| Female line (Taiwan) | -0.157 | 0.064 | -2.457 | 0.015 * |
| Second male line (Greece) | -0.199 | 0.063 | -3.165 | 0.002 ** |
| Second male line (Malaysia) | 0.003 | 0.063 | 0.052 | 0.958 |
| Second male line (Taiwan) | -0.151 | 0.064 | -2.364 | 0.019 * |
| Treatment (het-perfumed) | -0.071 | 0.037 | -1.946 | 0.054 |
| Female line (Greece) x second male line (Greece) | -0.014 | 0.105 | -0.129 | 0.897 |
| Female line (Malaysia) x second male line (Greece) | 0.049 | 0.110 | 0.443 | 0.658 |
| Female line (Taiwan) x second male line (Greece) | 0.093 | 0.110 | 0.841 | 0.402 |
| Female line (Greece) x second male line (Malaysia) | 0.203 | 0.110 | 1.849 | 0.066 |
| Female line (Malaysia) x second male line (Malaysia) | -0.005 | 0.110 | -0.047 | 0.963 |
| Female line (Taiwan) x second male line (Malaysia) | -0.044 | 0.110 | -0.397 | 0.692 |
| Female line (Greece) x second male line (Taiwan) | -0.202 | 0.110 | -1.840 | 0.068 |
| Female line (Malaysia) x second male line (Taiwan) | 0.141 | 0.111 | 1.278 | 0.203 |
| Female line (Taiwan) x second male line (Taiwan) | 0.129 | 0.111 | 1.162 | 0.247 |

| Table S5. Variation in female second-mating frequency depending on female line, second male line, and treatment group, as assessed in a beta-binomial generalized linear model. California females, California males, and conspecific-perfumed males are incorporated into the model as a baseline.  Alt text: A table containing the output of our beta-binomial generalized linear model. Our model includes significant terms for at least some components of female genotype and second male genotype, but no significant effects from perfuming treatment. | | | | |
| --- | --- | --- | --- | --- |
| Model component | **Estimate** | **Std. error** | ***z*** | ***p*** |
| Intercept | 1.123 | 0.160 | 7.021 | <0.001 *** |
| Female line (Greece) | -0.636 | 0.228 | -2.792 | 0.005 ** |
| Female line (Malaysia) | 0.333 | 0.265 | 1.258 | 0.209 |
| Female line (Taiwan) | -0.620 | 0.226 | -2.742 | 0.006 ** |
| Second male line (Greece) | -0.652 | 0.224 | -2.914 | 0.004 ** |
| Second male line (Malaysia) | -0.385 | 0.227 | -1.697 | 0.090 |
| Second male line (Taiwan) | 0.485 | 0.268 | 1.812 | 0.070 |
| Treatment (het-perfumed) | -0.038 | 0.138 | -0.275 | 0.783 |

| Table S6. Mean trait values and standard deviations for morphology data across all lines.  Alt text: A table containing the means and standard deviations of morphological traits across genotypes. | | | | | |
| --- | --- | --- | --- | --- | --- |
| Trait | **Line** | | | | |
|  | **GFP** | **California** | **Greece** | **Malaysia** | **Taiwan** |
| Female thorax length (mm) | 0.951  ± 0.045 | 0.882  ± 0.040 | 0.906  ± 0.030 | 0.835  ± 0.053 | 0.845  ± 0.022 |
| SR length (mm) | 2.382  ± 0.452 | 2.134  ± 0.172 | 2.059  ± 0.086 | 1.901  ± 0.139 | 2.061  ± 0.125 |
| ST area (mm^2^) | 0.005  ± 0.001 | 0.005  ± 0.000 | 0.005  ± 0.001 | 0.004  ± 0.001 | 0.005  ± 0.001 |
| Female total dry body mass (mg) | 0.383  ± 0.012 | 0.383  ± 0.024 | 0.339  ± 0.031 | 0.356  ± 0.084 | 0.411  ± 0.030 |
| FRT dry mass (mg) | 0.089  ± 0.012 | 0.095  ± 0.019 | 0.063  ± 0.012 | 0.082  ± 0.037 | 0.054  ± 0.028 |
| Male thorax length (mm) | 0.867  ± 0.035 | 0.789  ± 0.043 | 0.837  ± 0.046 | 0.764  ± 0.035 | 0.719  ± 0.026 |
| Testis length (mm) | 2.735  ± 0.108 | 2.858  ± 0.130 | 2.868  ± 0.175 | 2.342  ± 0.181 | 2.344  ± 0.143 |
| Sperm length (mm) | 1.713  ± 0.059 | 1.814  ± 0.033 | 1.609  ± 0.024 | 1.662  ± 0.022 | 1.763  ± 0.022 |
| Male total dry body mass (mg) | 0.251  ± 0.011 | 0.254  ± 0.026 | 0.234  ± 0.023 | 0.277  ± 0.045 | 0.292  ± 0.028 |
| MRT dry mass (mg) | 0.026  ± 0.003 | 0.025  ± 0.004 | 0.029  ± 0.004 | 0.028  ± 0.003 | 0.022  ± 0.002 |

| Table S7. Mean trait values and standard deviations for second male courtship effort (courtship interactions per minute), female second-mating proportion, and second male paternity (P2) across all line combinations. C indicates trait values for conspecific-perfumed males, while H indicates data for heterospecific-perfumed males.  Alt text: A table containing the means and standard deviations of courtship effort, second-mating proportion, and second male paternity across genotypes and perfuming treatments. | | | | | |
| --- | --- | --- | --- | --- | --- |
|  | | **Male line** | | | |
|  |  | **California** | **Greece** | **Malaysia** | **Taiwan** |
| Female line | **California** | *Courtship effort*  C: 1.537 ± 1.475  H: 1.05 ± 0.315 | *Courtship effort*  C: 0.281 ± 0.158  H: 0.263 ± 0.253 | *Courtship effort*  C: 0.555 ± 0.397  H: 0.347 ± 0.254 | *Courtship effort*  C: 0.395 ± 0.301  H: 0.367 ± 0.198 |
|  |  | *Second-mating proportion*  C: 1.000  H: 1.000 | *Second-mating proportion*  C: 0.857  H: 0.750 | *Second-mating proportion*  C: 0.857  H: 0.714 | *Second-mating proportion*  C: 1.000  H: 0.857 |
|  |  | *P2*  C: 0.777 ± 0.246  H: 0.799 ± 0.169 | *P2*  C: 0.440 ± 0.294  H: 0.454 ± 0.315 | *P2*  C: 0.418 ± 0.267  H: 0.504 ± 0.361 | *P2*  C: 0.458 ± 0.287  H: 0.533 ± 0.266 |
|  | **Greece** | *Courtship effort*  C: 1.078 ± 0.402  H: 0.963 ± 0.500 | *Courtship effort*  C: 0.633 ± 0.515  H: 0.265 ± 0.254 | *Courtship effort*  C: 1.074 ± 0.589  H: 0.662 ± 0.219 | *Courtship effort*  C: 0.370 ± 0.269  H: 0.247 ± 0.135 |
|  |  | *Second-mating proportion*  C: 0.625  H: 0.556 | *Second-mating proportion*  C: 0.385  H: 0.417 | *Second-mating proportion*  C: 0.889  H: 0.600 | *Second-mating proportion*  C: 0.556  H: 0.875 |
|  |  | *P2*  C: 0.969 ± 0.044  H: 0.913 ± 0.088 | *P2*  C: 0.874 ± 0.149  H: 0.448 ± 0.229 | *P2*  C: 0.519 ± 0.271  H: 0.816 ± 0.196 | *P2*  C: 0.900 ± 0.155  H: 0.986 ± 0.019 |
|  | **Malaysia** | *Courtship effort*  C: 0.922 ± 0.318  H: 0.670 ± 0.268 | *Courtship effort*  C: 0.514 ± 0.512  H: 0.455 ± 0.193 | *Courtship effort*  C: 0.490 ± 0.242  H: 0.776 ± 1.31 | *Courtship effort*  C: 0.732 ± 0.708  H: 0.518 ± 0.517 |
|  |  | *Second-mating proportion*  C: 0.857  H: 1.000 | *Second-mating proportion*  C: 1.000  H: 0.667 | *Second-mating proportion*  C: 0.615  H: 0.625 | *Second-mating proportion*  C: 1.000  H: 0.778 |
|  |  | *P2*  C: 0.852 ± 0.139  H: 0.842 ± 0.195 | *P2*  C: 0.498 ± 0.269  H: 0.501 ± 0.227 | *P2*  C: 0.411 ± 0.281  H: 0.465 ± 0.299 | *P2*  C: 0.691 ± 0.278  H: 0.871 ± 0.136 |
|  | **Taiwan** | *Courtship effort*  C: 0.582 ± 0.291  H: 0.602 ± 0.479 | *Courtship effort*  C: 0.338 ± 0.073  H: 0.295 ± 0.174 | *Courtship effort*  C: 0.428 ± 0.257  H: 0.337 ± 0.237 | *Courtship effort*  C: 0.458 ± 0.252  H: 0.343 ± 0.102 |
|  |  | *Second-mating proportion*  C: 0.625  H: 1.000 | *Second-mating proportion*  C: 0.357  H: 0.625 | *Second-mating proportion*  C: 0.636  H: 0.385 | *Second-mating proportion*  C: 0.667  H: 0.800 |
|  |  | *P2*  C: 0.915 ± 0.065  H: 0.817 ± 0.136 | *P2*  C: 0.367 ± 0.362  H: 0.610 ± 0.231 | *P2*  C: 0.264 ± 0.247  H: 0.206 ± 0.167 | *P2*  C: 0.759 ± 0.187  H: 0.641 ± 0.220 |

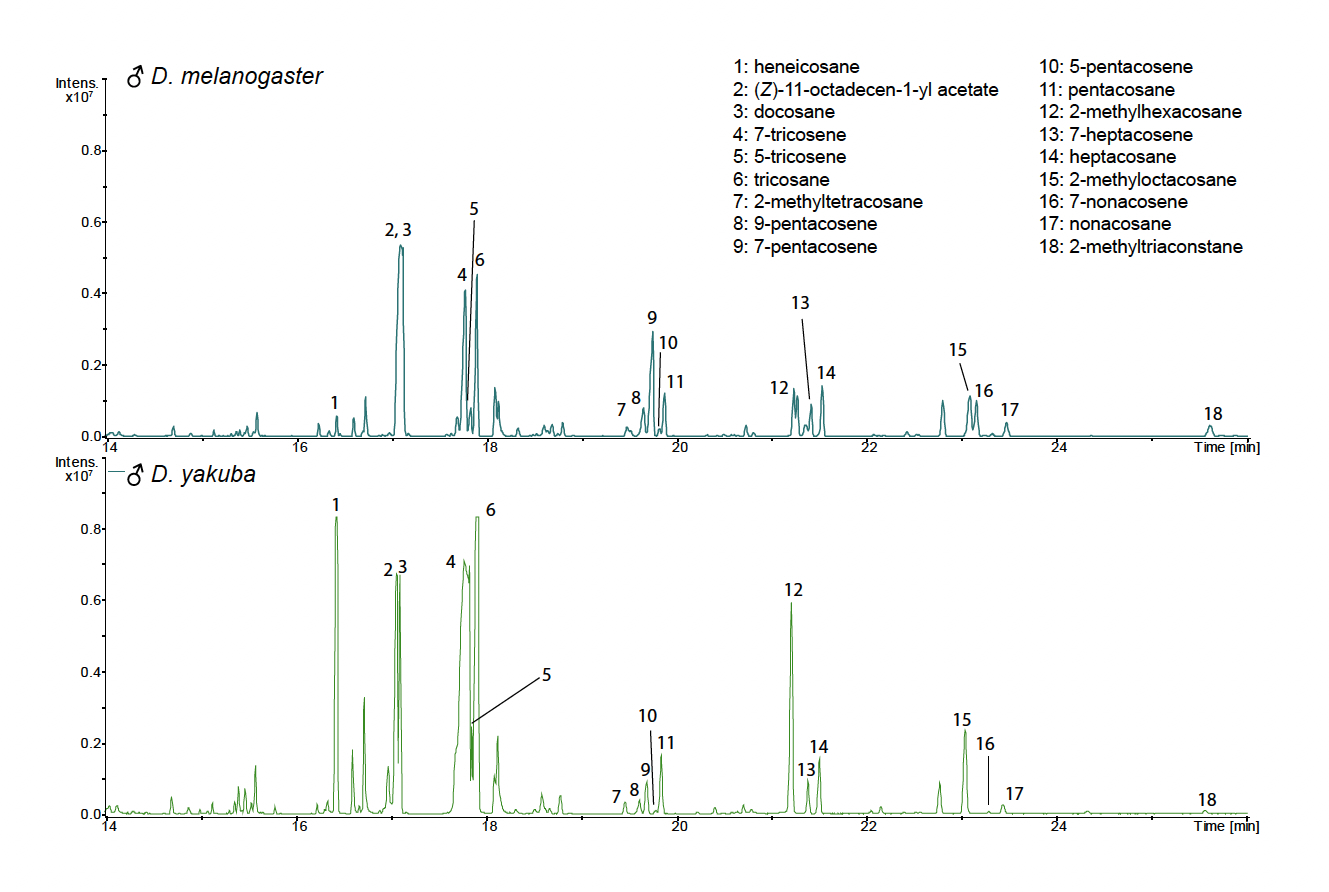

**Figure S1.** Representative gas chromatography mass spectrometry (GCMS) chromatograms of cuticular lipid extracts of male *D. melanogaster* and *D. yakuba*. Peaks corresponding to major cuticular compounds are labeled. Raw GCMS data from Khallaf et al. (2021) were downloaded and reanalyzed using Compass DataAnalysis software, V. 6.1 (Bruker Daltonics GmbH & Co., Germany).

**Alt text:** A graphical representation of GCMS data for *D. melanogaster* and *D. yakuba* males. A number of major cuticular compounds show substantial variation in quantity between the two species.

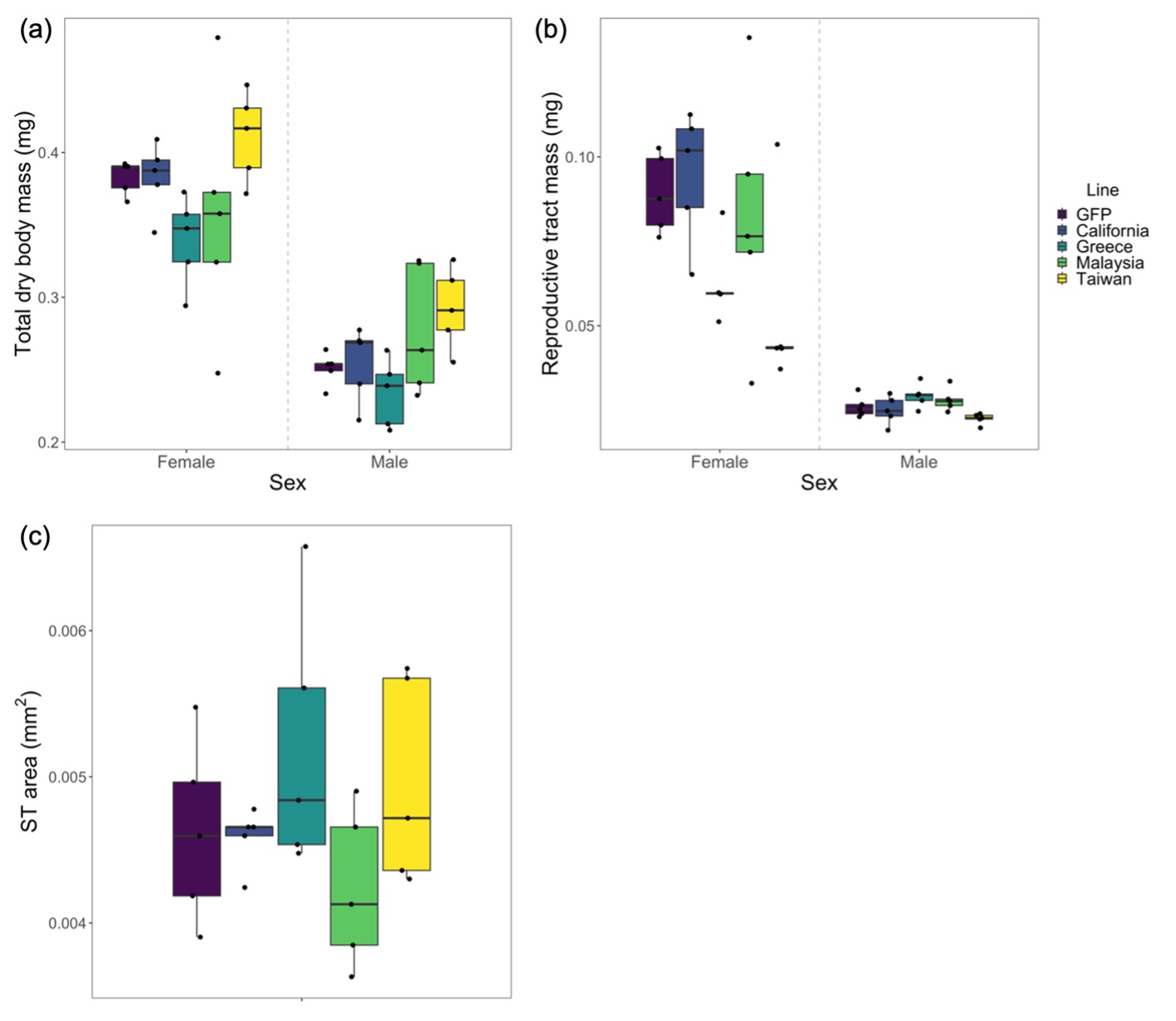

**Figure S2.** Variation in (a) total dry body mass in mg and (b) dry reproductive tract mass in mg in females and males across *D. melanogaster* lines (Table S3). Variation in (c) spermatheca area in mm^2^ is also shown across *D. melanogaster* female lines. Box plots show the median (central line) and first and third quartiles; whiskers extend to the smallest and largest values (not including outliers >1.5x the inter-quartile range).

**Alt text:** A graphical representation of variation in morphology across male and female *D. melanogaster* lines. Data are represented using box plots and individual data points.

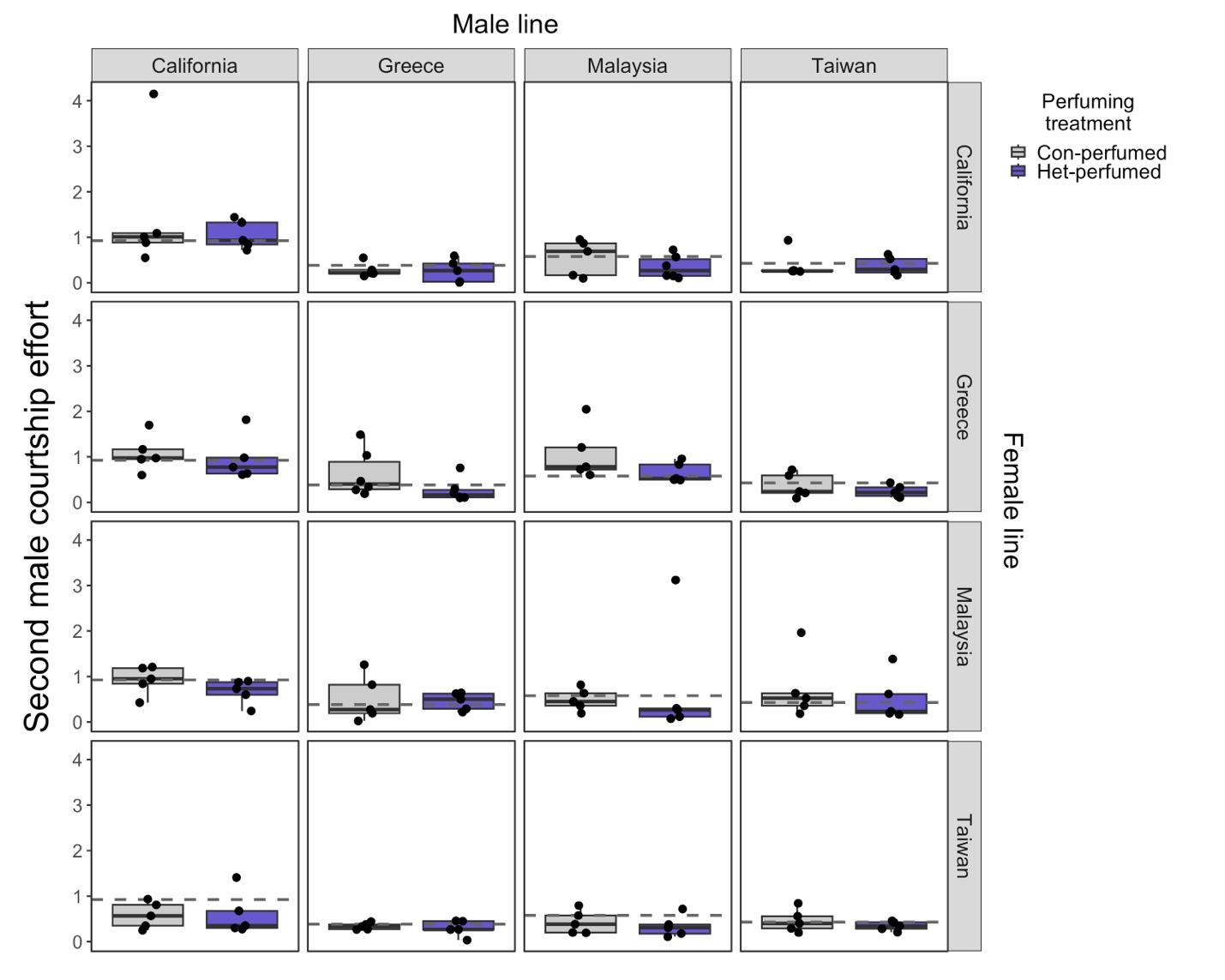

**Figure S3.** Courtship effort (courtship behaviors per minute) consistently differed between second male genotypes, but did not consistently vary by the female genotype being courted or by male perfuming status (Table 1, Table S4). Conspecific-perfumed males are shown in grey, while heterospecific-perfumed males are shown in purple. The dashed line indicates the average courtship effort for each male line. Box plots show the median (central line) and first and third quartiles; whiskers extend to the smallest and largest values (not including outliers >1.5x the inter-quartile range). This version includes two additional data points (both >2 courtship behaviors per minute) that were excluded from Figure 3.

**Alt text:** A matrix of graphs representing second male courtship effort across all possible male by female combinations, divided according to conspecific- or heterospecific-perfumed treatments. Data are represented using box plots and individual data points.

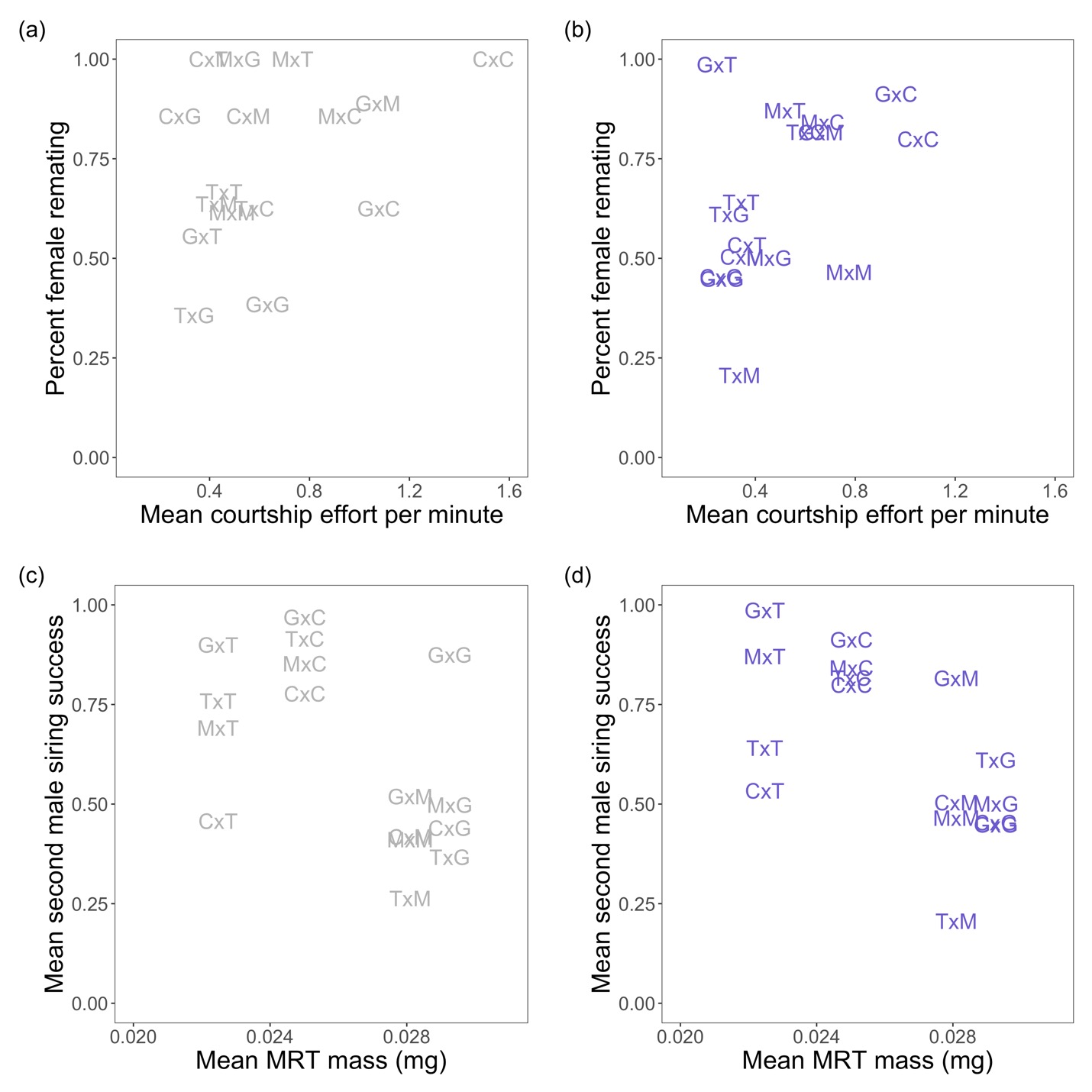

**Figure S4.** Courtship effort is not correlated with female remating rate when males are conspecific-perfumed (a) or heterospecific-perfumed (b). MRT mass is correlated with second male siring success in heterospecific-perfumed males (d), but not conspecific-perfumed males (c). Crosses are labeled as female x male genotype (G, Greece; T, Taiwan; C, California; and M, Malaysia).

**Alt text:** Scatterplots representing the correlation between datasets. Points are represented using the first letter of each genotype in that cross (for example, C x G is California female by Greece male). Data is divided between conspecific-perfumed and heterospecific-perfumed males.
